# Lipidomic profiling reveals age-dependent changes in complex plasma membrane lipids that regulate neural stem cell aging

**DOI:** 10.1101/2022.08.18.503095

**Authors:** Xiaoai Zhao, Xin Yan, Kévin Contrepois, Francesco Vallania, Mathew Ellenberger, Chloe M. Kashiwagi, Stephanie D. Gagnon, Cynthia J. Siebrand, Matias Cabruja, Gavin M. Traber, Andrew McKay, Daniel Hornburg, Purvesh Khatri, Michael P. Snyder, Richard N. Zare, Anne Brunet

**Affiliations:** Department of Genetics, Stanford University, CA, USA; Department of Chemistry, Stanford University, CA, USA; Stanford Center for Biomedical Informatics Research, Stanford University, CA, USA; Glenn Laboratories for the Biology of Aging, Stanford University, CA, USA; Department of Chemistry, Texas A&M University, TX, USA

## Abstract

The aging brain exhibits a decline in the regenerative populations of neural stem cells (NSCs), which may underlie age-associated defects in sensory and cognitive functions^1–6^. While mechanisms that restore old NSC function have started to be identified^7–23^, the role of lipids – especially complex lipids – in NSC aging remains largely unclear. Using lipidomic profiling by mass spectrometry, we identify age-related lipidomic signatures in young and old quiescent NSCs *in vitro* and *in vivo*. These analyses reveal drastic changes in several complex membrane lipid classes, including phospholipids and sphingolipids in old NSCs. Moreover, poly-unsaturated fatty acids (PUFAs) strikingly increase across complex lipid classes in quiescent NSCs during aging. Age-related changes in complex lipid levels and side chain composition are largely occurring in plasma membrane lipids, as revealed by lipidomic profiling of isolated plasma membrane vesicles. Experimentally, we find that aging is accompanied by modifications in plasma membrane biophysical properties, with a decrease in plasma membrane order in old quiescent NSCs *in vitro* and *in vivo*. To determine the functional role of plasma membrane lipids in aging NSCs, we performed genetic and supplementation studies. Knockout of *Mboat2*, which encodes a phospholipid acyltransferase, exacerbates age-related lipidomic changes in old quiescent NSCs and impedes their ability to activate. As *Mboat2* expression declines with age, *Mboat2* deficiency may drive NSC decline during aging. Interestingly, supplementation of plasma membrane lipids derived from young NSCs boosts the ability of old quiescent NSCs to activate. Our work could lead to lipid-based strategies for restoring the regenerative potential of NSCs in old individuals, which has important implications for countering brain decline during aging.

---

Lipids are as diverse and complex as proteins. Yet our understanding of lipid function is lagging. Complex lipids – lipids with more than one chemical moieties – are key components of cellular membranes (plasma and organelles), thereby serving critical barrier function as well as regulating organellar homeostasis and signal transduction^24–27^. In the brain, complex lipids are particularly abundant, and their dysregulation has been observed in brain aging and neurodegenerative diseases, such as Alzheimer’s disease^28–36^. But the role of complex lipids in cells in the brain – and other tissues – during aging is largely unknown.

The adult mammalian brain contains populations of neural stem cells (NSCs) that can generate neurons, astrocytes, and oligodendrocytes^1, 2, 37–41^. NSCs are organized in two main niches – the dentate gyrus of the hippocampus and the subventricular zone^39, 40, 42–45^. During aging, the ability of NSCs to transition from quiescence to activation declines^15, 46–50^, and this decline could underlie defects in aspects of cognitive and sensory function, as well as impaired injury repair in old individuals^51–54^. Understanding how to maintain NSC regenerative potential during aging could identify strategies to counter age-related decline and facilitate repair after injury in the central nervous system. While lipid energy metabolism has been shown to be important for NSCs and other stem cells^55–63^, the contribution of complex lipids – notably membrane lipids – in stem cell function during aging has not been studied. More generally, a systematic examination of the global lipidome of cells during aging is still missing.

## Lipidomics reveals changes in complex membrane lipids in old quiescent NSCs

To unbiasedly determine the age-related changes in complex lipids, we performed untargeted lipidomic profiling of NSCs from young and old mice in culture and *in vivo*. Using liquid chromatography followed by tandem mass spectrometry (LC-MS/MS), we profiled quiescent NSCs (qNSCs) and activated NSCs (aNSCs) primary cultures from young (3-5 month-old) and old (20-22 month-old) mice (Fig. 1a, Supplementary Table 1). We identified 373 unique lipids, covering all major lipid categories (Extended Data Fig. 1a). Principal component analysis (PCA) on all lipids easily separated NSCs by cell state (quiescent vs. activated) and by age for qNSCs (Fig. 1b). Complex lipid species involved in membranes, such as phosphatidylcholine (PC), phosphatidylethanolamine (PE), phosphatidylinositol (PI), and sphingomyelin (SM), showed strong changes (increase or decrease) with age, notably in quiescent NSCs (Fig. 1c,d). For example, species with age-related changes in phosphatidylserines (PS) and the majority of PI showed an increase with age, whereas ether lipid species of phosphatidylcholines (PC) and phosphatidylethanolamines (PE) decreased with age (Fig. 1c,d). Cholesterol, another major membrane constituent, also increased with age in qNSCs (Fig. 1c,d). In contrast, lipids involved in energy storage, such as diacylglycerol (DG) and triacylglycerol (TG), showed fewer changes with age (Fig. 1c,d). These results were corroborated in a second independent experiment (Extended Data Fig. 1b-e, Supplementary Table 2) and a third independent experiment using the targeted lipidomic platform Lipidyzer^64^ (Extended Data Fig. 1f-h, Supplementary Table 3). Thus, lipidomic profiling in NSC cultures uncovers large changes in complex membrane lipids with age, notably in quiescent NSCs.

**Figure 1.**
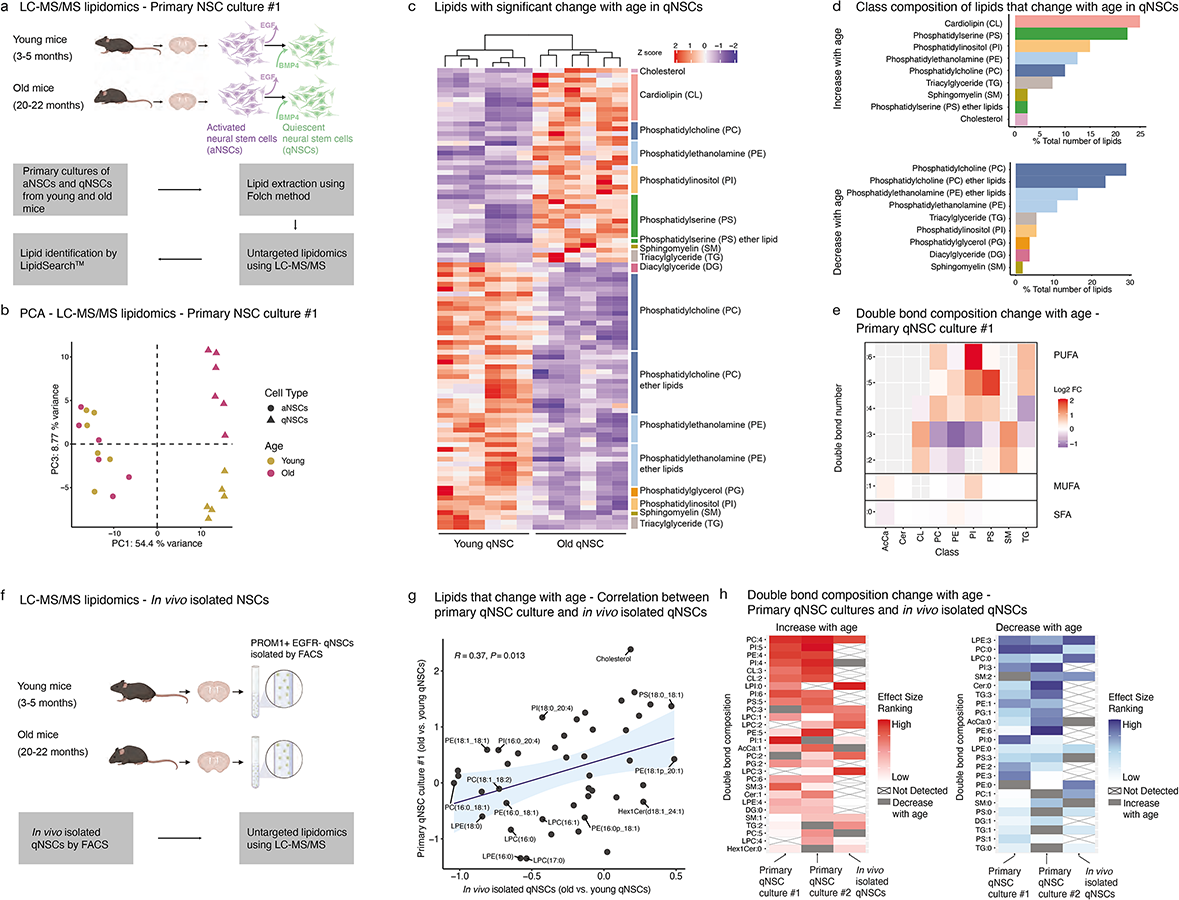
Lipidomics on young and old neural stem cells *in vitro* and *in vivo* identifies membrane lipid and side chain composition changes with age. **a,** Overview of lipidomic study *in vitro* (for *in vivo* design, see **f**). Primary NSC cultures were established from 6 young (3-5 months) and 6 old (20-22 months) C57BL/6JN male mice. Each culture was from an individual mouse. In each culture, activated NSCs (aNSCs) were expanded. In parallel, quiescent NSCs (qNSCs) were generated by the addition of bone morphogenetic protein 4 (BMP4) and the removal of epidermal growth factor (EGF). Lipids from activated and quiescent NSC cultures were extracted using a modified Folch method and analyzed using an untargeted lipidomic platform with liquid chromatography followed by tandem mass spectrometry (LC-MS/MS). Lipids were identified by LipidSearch^TM^ for downstream analysis. **b,** Principal component analysis (PCA) on log2-transformed normalized intensity of all lipids for primary NSC culture #1 (see Extended Data Fig. 1c for data with primary NSC culture #2). Each symbol represents an individual primary aNSC culture (dot) or qNSC culture (triangle) established from individual young (golden) or old (pink) mice. **c,** Heatmap with clustering on the Z score of normalized lipid intensity for 95 lipids with significant change with age (FDR<0.05) in qNSCs for primary qNSC culture #1 (see Extended Data Fig. 1d for correlation with primary qNSC culture #2). Lipids were separated into 2 categories (increase with age (top) or decrease with age (bottom)) and then grouped by lipid classes. **d,** Class composition of lipids that significantly change with age (age-related lipids) in qNSCs for primary qNSC culture #1 (see Extended Data Fig. 1e for primary qNSC culture #2). Number of lipids in each lipid class as a percentage of the total number of lipids that show significant increase with age (top panel) or decrease with age (bottom panel). **e,** Heatmap showing double bond composition change with age between young and old qNSCs for side chain analysis for primary qNSC culture #1 (see Extended Data Fig. 1i for primary qNSC culture #2). Log2 fold change in abundance of lipids with side chain containing between 0-6 double bonds (rows) in different lipid classes (columns) are shown. Abbreviations defined in panel d, except for AcCa: acylcarnitine and Cer: ceramide. **f,** Schematic of lipidomic profiling on qNSCs *in vivo*. Quiescent NSCs were freshly isolated from the brain of young (3-5 months) or old (20-22 months) C57BL/6JN male mice using fluorescence-activated cell sorting (FACS) based on the positive staining of Prominin-1 (PROM1+) and negative staining of epidermal growth factor receptor (EGFR-). Isolated cells were then subjected to untargeted lipidomic analysis using LC-MS/MS. **g,** Correlation between lipid changes with age in qNSCs from primary culture (primary qNSC culture #1) and *in vivo*. Pearson correlation performed on effect size of changes with age of individual lipids (black dots) from each dataset. Linear regression line (black line) with 95% confidence interval (blue shade). Membrane lipids in the top or bottom 20 percentile of either dataset are annotated. **h,** Double bond composition change with age in qNSCs from primary cultures #1 and #2 and *in vivo*. Increase with age (left panel) or decrease with age (right panel). The effect size of double bond composition change with age was calculated and used to generate a percentile ranking for each dataset. Consistent double bond composition features that were detected in at least 2 out of the 3 datasets were then sorted based on the combined percentile ranking across all 3 studies. Heatmap was plotted using the percentile ranking of double bond features in each lipidomic dataset. Double bond features not detected are crossed out. Gray indicates double bond features with opposite age-related changes.

The composition and unsaturation level of fatty acyl side chains in each lipid class is critical for lipid biophysical properties, especially at membranes^24, 26^. In qNSCs, the abundance of poly-unsaturated fatty acids (PUFAs) side chains increased in old qNSCs across many lipid classes, including membrane lipids (Fig. 1e). Notably, highly-unsaturated PUFAs (4 or more double bonds) drastically increased with age (Fig. 1e, Extended Data Fig. 1i). In contrast, the abundance of saturated fatty acids (SFAs) and mono-unsaturated fatty acids (MUFAs) side chains remained mostly unchanged with age (Fig. 1e). Hence, quiescent NSCs in primary culture exhibit age-related accumulation of highly-unsaturated PUFA side chains in complex membrane lipids.

To determine whether the changes in primary NSC cultures are recapitulated *in vivo*, we performed lipidomic profiling on freshly isolated qNSCs from the subventricular zone (SVZ) neurogenic niche of young and old mouse brains^14, 15, 65^ (Fig. 1f, Extended Data Fig. 8a, Supplementary Table 4). Untargeted lipidomics using LC-MS/MS on qNSCs freshly isolated by fluorescence-activated cell sorting (FACS) identified 128 lipids in qNSCs, including major lipid classes identified in primary cultures (Extended Data Fig. 2a). Data from *in vivo* isolated qNSCs showed some separation with age by PCA (Extended Data Fig. 2b). The age-related changes in lipid species in freshly isolated qNSCs correlated at least in part with changes in primary cultures (Fig. 1g), suggesting consistent differences in lipidome with age *in vitro* and *in vivo*. In addition, side chain composition analysis in freshly isolated qNSCs also showed several unsaturation level changes with age similar to those observed in primary qNSC cultures, including increased abundance of PC containing 4-double bond (PC:4) side chains and decreased abundance of lysophosphatidylethanolamine (LPE) containing 3-double bond side chains (LPE:3) (Fig. 1h, Extended Data Fig. 2c). Collectively, these lipidomic results indicate that quiescent NSCs *in vitro* and *in vivo* exhibit large changes in complex membrane lipids and accumulate highly unsaturated PUFA side chains during aging.

## Spatial lipidomic profiling also shows changes in complex lipids in quiescent NSCs in old neurogenic niches

We next asked how lipids are modulated *in situ* in old neurogenic niches. To identify lipids *in situ,* we used desorption electrospray ionization mass spectrometry imaging (DESI-MSI) on whole mount sections from SVZ niches (Fig. 2a, Supplementary Table 5). DESI-MSI enables simultaneous *in situ* detection of lipids and metabolites while preserving tissue integrity^66, 67^ (Fig. 2a). Given DESI-MSI’s spatial resolution (200μm), each individual mass spectrometry spectrum of the SVZ niche encompasses metabolites of several cell types in a mixed population (Fig. 2a). To obtain cell type-specific lipidomic profiles, we aligned DESI-MSI spectra data with immunofluorescence image of cell type-specific markers (GFAP+Ki67-for qNSCs, GFAP+Ki67+ for aNSCs, and GFAP-Ki67+ for neural progenitor cells, NPCs) (Fig. 2a-c). We then performed computational deconvolution on DESI-MSI spectra with cell type composition data using linear regression^68, 69^ to obtain lipidomic profiles of qNSCs, aNSCs and NPCs for each individual animal (Fig. 2a). This deconvolution method yielded accurate estimates of cell type-specific profiles from a reconstituted *in silico* mixture lipidomic dataset (see Methods) (Extended Data Fig. 3a,b). Lipidomic profiles from deconvolved *in situ* qNSCs showed some separation with age by PCA (Fig. 2d). We then used a supervised approach – sparse partial least square discriminant analysis (sPLS-DA) – to select for features (lipids and/or metabolites) that showed specific changes between cell types and by age (see Methods) (Extended Data Fig. 3c,d). Using this approach, we identified lipids and/or metabolites that exhibited similar changes with age across all three cell types (qNSCs, aNSCs, and NPCs) (Extended Data Fig. 3e). In addition, we also identified mass spectra (corresponding to lipids and/or metabolites) with cell type-specific changes with age (Extended Data Fig. 3f).

**Figure 2.**
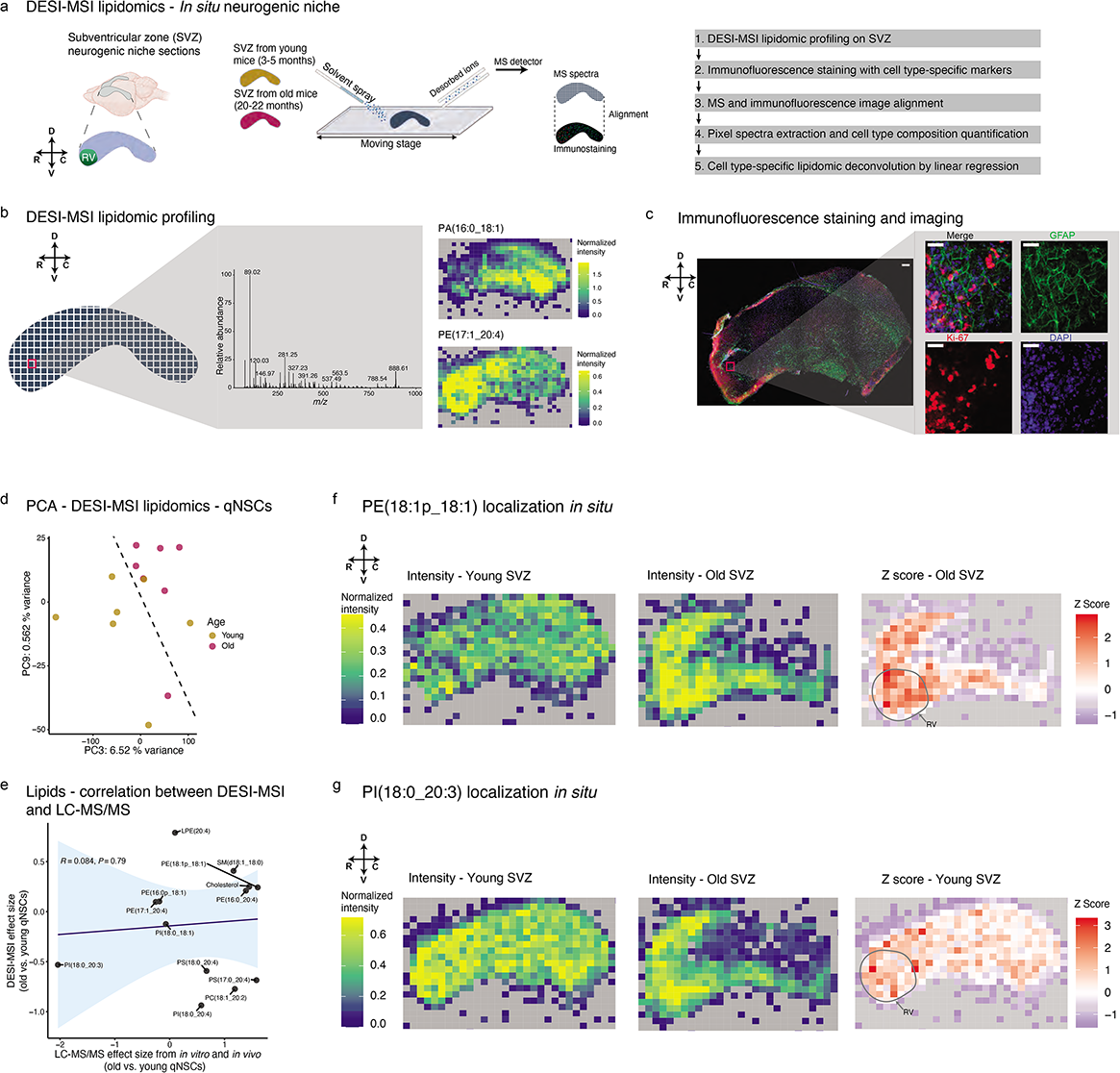
*In situ* lipidomic profiling on neural stem cells from the subventricular zone neurogenic niche of young and old mice shows membrane lipids change with aging. **a,** Overview of lipidomic study *in situ* using desorption electrospray ionization mass spectrometry imaging (DESI-MSI). Subventricular zone (SVZ) neurogenic niche sections of whole mount preparation from 7 young (3-5 months) and 7 old (20-22 months) C57BL/6JN male mice were placed onto a moving stage. Solvent spray was introduced with a 200μm nozzle, and desorbed ions across the entire SVZ neurogenic niche were detected by mass spectrometry (MS) in an array. Arrows indicate orientation axes, with R: rostral, C: caudal, D: dorsal, and V: ventral. RV: rostral-ventral region. Sections are analyzed by DESI-MSI and then stained by immunofluorescence staining for cell type markers. The mass spectra and immunofluorescence images are then aligned, and deconvolution for cell type-specific lipidomic profile is done computationally using a linear regression method on each individual animal. **b,** Left panel: representative mass spectra from an individual DESI-MSI scan showing the abundance of metabolites (including lipids) with *m/z* between 50-1000. Each individual pixel represents a separate scan by DESI-MSI. Right panel: representative DESI-MSI image showing the tissue distribution of PA(16:0_18:1) (*m/z*: 673.5) (top) and PE(17:1_20:4) (*m/z*: 750.5) (bottom) across the SVZ neurogenic niche. Heatmap generated based on normalized lipid intensity. **c,** Left panel: immunofluorescence staining of SVZ section from whole mount preparation, post-lipidomic profiling by DESI-MSI. Staining with GFAP (NSC and astrocyte marker, green), Ki67 (proliferation marker, red) and DAPI (nuclei, blue). Scale bar: 200μm. Right panel: enlarged immunofluorescence images covering the area of one individual DESI-MSI scan. Same set of markers as left panel. Scale bar: 50μm. **d,** Principal component analysis (PCA) on estimated intensity of all lipids and other metabolites in young and old qNSCs *in situ*. Each point represents a deconvolved qNSC lipidomic profile from individual young (3-5 months, golden) or old (20-22 months, pink) C57BL/6JN male mouse. A computed decision boundary indicating the separation between young and old samples (see Methods) was added to the PCA plot (dashed line). **e,** Correlation analysis on lipid changes with age in qNSCs between *in situ* lipidomics by DESI-MSI, and *in vitro* and *in vivo* lipidomics by LC-MS/MS. Pearson correlation was performed on the effect size of changes with age of individual lipids (black dots) in DESI-MSI dataset and the summary effect size of changes with age from meta-analysis (see Methods) of lipids from 2 primary quiescent neural stem cell (qNSC) lipidomic datasets and *in vivo* isolated qNSC lipidomic dataset. Linear regression line (black line) with 95% confidence interval (blue shade). **f, g,** Lipid distribution *in situ*. Heatmap with normalized intensity of PE(18:1p_18:1) (**f**) and PI(18:0_20:3) (**g**) in a representative young (left) and old (middle) SVZ is shown. Z score heatmap (right) is shown in the old SVZ (**f**) and the young SVZ (**g**). RV region of the SVZ is indicated with gray circle.

To validate the specific lipids that changed with age *in situ* in DESI-MSI datasets, we performed an additional tandem mass spectrometry step (Extended Data Fig. 4). We focused on lipids identified in qNSCs *in vitro* and *in vivo* (see Fig. 1). Overall, 7 out of 13 complex membrane lipids showed consistent changes in old qNSCs by DESI-MSI *in situ,* and lipidomic studies on qNSCs *in vitro* and *in vivo* (correlation coefficient *R* = 0.084, *P* value = 0.79), including PI(18:0_20:3), cholesterol, and PE(16:0_20:4) (Fig. 2e). Hence, complex membrane lipids change with age *in situ* in neurogenic niches.

We next examined the spatial pattern of membrane lipids that show age-related changes. We focused on PE(18:1p_18:1) and PI(18:0_20:3), which show an increase or decrease, respectively, in qNSCs with age (Fig. 2f, g). While these membrane lipids were both detected across the SVZ, the most pronounced changes with age (Z scores) were in the rostral-ventral (RV) region of the SVZ (Fig. 2f-g). As the RV region has high qNSC density^70–72^, these data are consistent with the stronger changes in membrane lipids in qNSCs during aging. Thus, lipidomic profiling by DESI-MSI reveals age-related lipidomic changes in qNSCs *in situ* in the old neurogenic niche, including membrane lipids.

## Plasma membrane lipids and biophysical properties are affected during aging

Membrane lipids are major components of both plasma and organelle membranes (endoplasmic reticulum, Golgi apparatus, lysosome, and mitochondrion)^73, 74^. To assess the organelles and cellular compartments that are the most affected during aging in qNSCs, we performed lipid ontology enrichment analysis^75^. This analysis revealed that lipids from the plasma membrane were significantly enriched in old qNSCs (Fig. 3a). In contrast, lipids from endoplasmic reticulum were depleted in old qNSCs, and lipids from other organelles did not significantly change with age (Fig. 3a).

**Figure 3.**
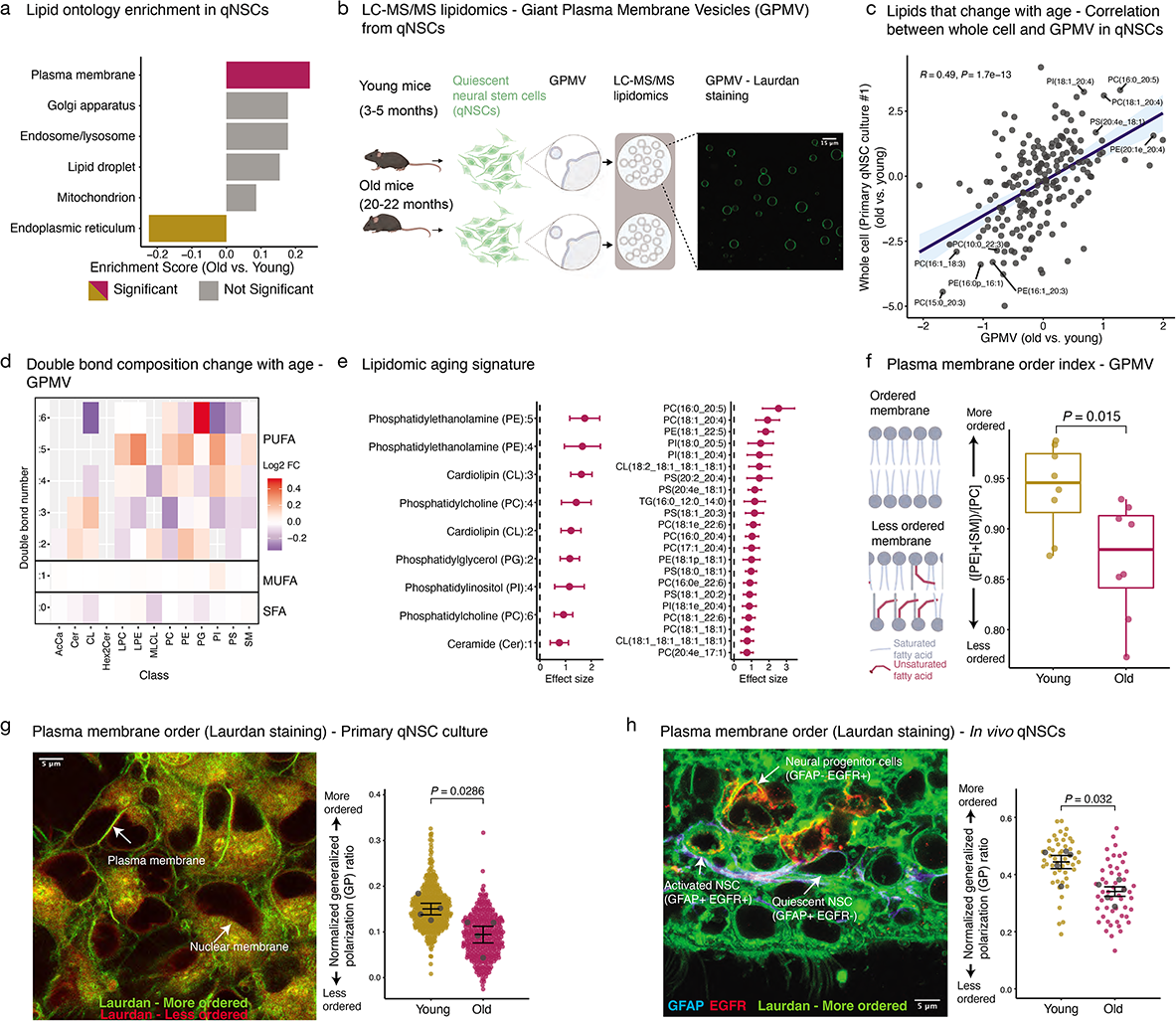
Plasma membrane lipids and biophysical properties change with age in quiescent neural stem cells. **a,** Lipid ontology analysis for different cellular compartments on lipids that change with age in qNSCs (lipidomic dataset from primary qNSC culture #1). Age-related enrichment score for each cellular compartment is presented. Positive enrichment score: enriched in old qNSCs. Negative enrichment score: enriched in young qNSCs. Significant terms (FDR<0.05) are shown in golden/pink color. Terms without statistical significance are shown in gray. **b,** Giant plasma membrane vesicles (GPMVs) were generated from quiescent neural stem cell (qNSC) primary cultures from 8 young (3-5 months) and 8 old (20-22 months) C57BL/6JN male mice, and were subjected to lipidomic analysis using LC-MS/MS. Confocal image showing isolated GPMVs visualized by staining with Laurdan (a dye that incorporates into membranes). **c,** Correlation between lipid changes in qNSCs with age between primary qNSC cultures (whole cell) and GPMVs from primary qNSC cultures. Pearson correlation performed on effect sizes of changes with age of individual lipids (black dots) calculated from each dataset. Linear regression line (black line) with 95% confidence interval (blue shade). Lipids in the top or bottom 10 percentile in both datasets are annotated. **d,** Heatmap showing double bond composition change with age in GPMV lipids from qNSCs, based on side chain composition analysis. Log2 fold change in abundance of lipids with side chain containing between 0-6 double bonds (rows) in different lipid classes (columns) are shown. Abbreviations defined in Figure 1, except for MLCL: monolysocardiolipin and Hex2Cer: dihexosylceramides. **e,** Lipidomic aging signature from meta-analysis (see Methods) on 2 primary qNSC culture lipidomic datasets, *in vivo* isolated qNSC lipidomic dataset, and qNSC GPMV lipidomic dataset. Dot plot showing the mean summary effect size +/-SEM of side chain unsaturation features (left) and individual lipid features (right) that exhibit a higher abundance with age. **f,** Plasma membrane order index calculated on GPMV lipids from young and old qNSCs. Plasma membrane order index was calculated as the ratio between the total concentration of phosphatidylethanolamine (PE) plus sphingomyelin (SM), and the total concentration of phosphatidylcholine (PC). Each dot represents GPMV lipids obtained from an individual primary qNSC culture. Results are from 1 independent experiment, n=8 independent primary qNSC cultures. *P*-value from Wilcoxon rank-sum test. **g,** Plasma membrane order assessed *in vitro* by Laurdan staining on young and old qNSCs. Left: representative Laurdan staining on primary qNSC cultures from 4 young (3-5 months) and 4 old (20-22 months) mice. Image from Laurdan rigid wavelength (more ordered, green) overlaid with Laurdan fluid wavelength (less ordered, red). Right: plasma membrane order quantified by generalized polarization ratio (GP ratio) from Laurdan staining of young and old qNSCs. SuperPlots showing data from individual cells (golden/pink dots), as well as the mean of each primary culture from individual mice (gray dots) +/-SEM. Results from one independent experiment, n=4 primary cultures. *P*-value from Wilcoxon rank-sum test. Results from a second independent experiment are in Supplementary Table 9. **h,** Plasma membrane order assessed *in vivo* by Laurdan staining of qNSCs in brain sections from 5 young (3-5 months) and 5 old (20-22 months) mice. Left: representative Laurdan staining of *in vivo* qNSCs from a brain section. Image from rigid wavelength (more ordered, green) overlaid with GFAP (NSC and astrocyte marker, blue) and EGFR (proliferation marker, red) staining. Right: plasma membrane order quantified by generalized polarization ratio (GP ratio) from Laurdan staining of qNSCs from 5 young (3-5 months) and 5 old (20-22 months) male mice. SuperPlots showing data from individual cells (golden/pink dots), as well as the mean of each mouse brain (gray dots) +/-SEM. Results from one independent experiment, n=5 mice. *P*-value from Wilcoxon rank-sum test. Result from a second independent experiment are in Supplementary Table 9.

To experimentally assess the age-related changes in plasma membrane lipids, we isolated plasma membrane from quiescent NSCs by generating giant plasma membrane vesicles (GPMVs) (Fig. 3b). GPMVs do not contain organellar lipids and represent mostly plasma membrane lipids in cells^76, 77^. We verified that GPMVs from qNSCs were devoid of organelles and other cellular compartments by western blot (Extended Data Fig. 5a, 8c) and that they were enriched for “Plasma membrane” lipids in lipidomic analysis (Extended Data Fig. 5b). Untargeted lipidomics on GPMVs derived from young and old qNSCs revealed that there was a strong positive correlation between the age-related changes in the lipidomes of whole cell extract and GPMVs (Fig. 3c, Supplementary Table 6). PUFAs – especially highly unsaturated PUFAs – accumulated in the side chains of the GPMV lipids from old qNSCs (Fig. 3d, Extended Data Fig. 5d). Thus, lipid changes in old quiescent NSCs mostly occur at the level of the plasma membrane.

We next generated “aging lipidomic signatures” by performing a meta-analysis on all of our lipidomics datasets from young and old qNSCs (see Methods). We identified 2 sets of signatures by assessing either the side chain unsaturation level (double bond composition) or individual lipid concentration change with age (Fig. 3e). The most robust aging lipid signature included 9 specific side chain unsaturation features and 22 individual lipid species, mostly plasma membrane lipids (Fig. 3e). Hence, changes in plasma membrane lipids (and other lipids) with age could be used to distinguish a youthful lipidomic profile from an older one (see Fig. 4 below).

**Figure 4.**
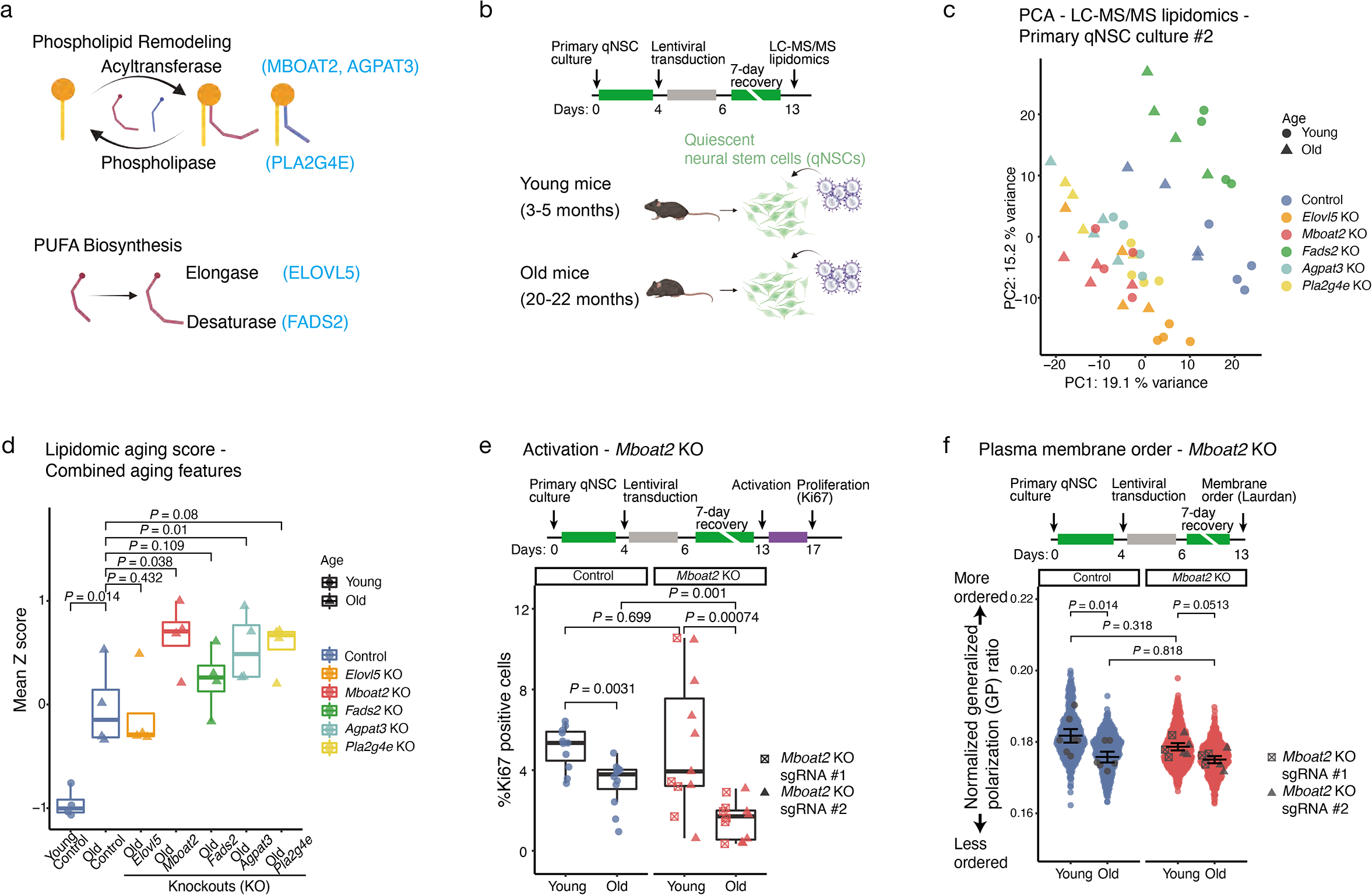
Knocking out the *Mboat2* acyltransferase gene exacerbates the age-related lipidomic and functional defects of old quiescent neural stem cells. **a,** Overview of the categories of enzymes involved in phospholipid remodeling pathway and poly-unsaturated fatty acid (PUFA) biosynthesis pathway. Genes selected for genetic knockout (KO) are highlighted in blue. **b,** Experimental setup. Primary NSC cultures from young (3-5 months) and old (20-22 months) mice were induced into quiescence for 4 days and then transduced with lentiviruses for CRISPR/Cas9 knockout of individual target genes for 2 consecutive days. Cells were then allowed a recovery period of 7 days before being subjected to untargeted lipidomic analysis by LC-MS/MS. **c,** Principal component analysis (PCA) on log2-transformed normalized values of all lipids. For lipids that have internal standard for their respective lipid classes, molar concentrations were used as values, for lipid classes that do not have internal standard, normalized intensity were used as values. Each symbol represents an individual primary qNSC culture established from young (dot) or old (triangle) mice. Cultures were either transduced with a control lentivirus with a non-targeting sgRNA (blue), or sgRNA targeting *Elovl5* (orange), *Mboat2* (red), *Fads2* (green), *Agpat3* (turquoise) or *Pla2g4e* (yellow). **d,** Lipidomic aging score calculated using combined aging signatures (side chain unsaturation and individual lipid) from young (blue, circle) and old control (blue, triangle) samples and old samples with *Elovl5* knockout (KO) (orange, triangle), *Mboat2* KO (red, triangle), *Fads2* KO (green, triangle), *Agpat3* KO (turquoise, triangle) or *Pla2g42* KO (yellow, triangle). Box and whisker plot from 1 independent experiment, n=4 primary cultures. Horizontal bar: median. Whisker: Distance between the first/third quartile to the highest/lowest value, respectively, or 1.5 times the distance between the first/third quartile, whichever is smaller. Each data point represents a primary culture from an individual mouse. *P*-values between old KO samples were from paired Welch’s *t*-test with multiple hypothesis correction by FDR. Normality of the data was confirmed for use of *t*-test. *P*-value between old and young control samples were from unpaired Welch’s *t*-test (young and old samples were not paired). Normality of the data was confirmed for use of *t*-test. **e,** Activation (proliferation) ability of young and old qNSCs with *Mboat2* KO. Primary qNSC cultures from young (3-5 months) and old (20-22 months) mice were transduced with control lentivirus (sgRNA targeting an unannotated region of the genome) or with sgRNAs targeting *Mboat2* (2 independent sgRNAs) for 2 consecutive days. After a 7-day recovery period, qNSCs were switched to proliferative NSC media for 4 days. The percentage of young and old control qNSCs (blue) and qNSCs with *Mboat2* KO (red) that successfully activated was assessed by intracellular FACS analysis for Ki67, a proliferation marker. Box and whisker plot from 4 independent experiments, n=11-12 primary cultures. Horizontal bar: median. Whisker: Distance between the first/third quartile to the highest/lowest value, respectively, or 1.5 times the distance between the first/third quartile, whichever is smaller. Each dot represents a primary culture from an individual mouse. *P*-values from Wilcoxon rank-sum test. Squares: sgRNA #1, Triangles: sgRNA #2. **f,** Plasma membrane order as assessed by Laurdan *in vitro* on young and old qNSCs with *Mboat2* knockout. Primary qNSC cultures from young (3-5 months) and old (20-22 months) mice were transduced with control lentivirus (sgRNA targeting an unannotated region of the genome) or with an sgRNA targeting *Mboat2* (2 independent sgRNAs) for 2 consecutive days. Plasma membrane order was assessed at the end of a 7-day recovery period following genetic KO. Plasma membrane order was quantified by generalized polarization ratio (GP ratio) from Laurdan staining on young and old control (blue) or *Mboat2* (red) KO qNSCs. SuperPlots showing data from individual cells (colored dots), as well as the mean of each primary culture (gray dots) +/-SEM. Results from 3 independent experiments, n=6-7 primary cultures. *P*-values from Wilcoxon rank-sum test.

Age-associated changes in lipids can impact the biophysical properties of plasma membranes, including plasma membrane order (i.e. rigidity or fluidity)^27, 78^. We used the lipid class composition from GPMVs to predict how membrane order is affected by age in qNSCs. As the abundance of specific phospholipid classes^79, 80^ is associated with different plasma membrane order, we calculated a “phospholipid order index”^81^ for GPMVs isolated from young and old qNSCs. To this end, we assessed the ratio between the concentration of phosphatidylethanolamine and sphingomyelin combined and phosphatidylcholine ([PE]+[SM])/[PC]^81^ in GPMVs (Fig. 3f). Plasma membranes derived from old qNSCs had a lower phospholipid order index – associated with lower plasma membrane order – compared to young qNSCs (Fig. 3f). In addition, plasma membranes from old qNSCs also had high phospholipid unsaturation^82^ and depletion in cholesterol (Extended Data Fig. 5c,d)^25, 80^, both of which are linked to lower plasma membrane order.

To experimentally test these predictions, we used the polarity-sensitive dye Laurdan to assess membrane order in primary NSC cultures and brain sections *in vivo*^83–85^. Ratiometric quantification of Laurdan at the plasma membrane provides a measure for membrane order (green: more ordered; red: less ordered). Quantification of Laurdan staining revealed that plasma membranes from old qNSC cultures had a lower order compared to their young counterparts (Fig. 3g), as predicted by the phospholipid order index. As an independent approach to assess plasma membrane order, we used fluorescence lifetime imaging (FLIM) to quantify fluorescence decay time – a metric that increases when membrane order is higher^86, 87^. FLIM analysis confirmed that qNSCs have decreased plasma membrane order with age (Extended Data Fig. 5e). Furthermore, Laurdan staining on brain sections from young and old mice showed that qNSCs also exhibit decreased plasma membrane order in old animals *in vivo* (Fig. 3h), both around the cell body and at the apical process (Fig. 3h, Extended Data Fig. 5f). Together, these results indicate that age-related lipidomic changes in qNSCs occur largely at the plasma membrane and are associated with a key biophysical property – a less ordered (more fluid) plasma membrane.

## Knockout of the phospholipid acyltransferase MBOAT2 exacerbates NSC aging

We asked whether pathways involved in the remodeling or biosynthesis of lipids that change with age in qNSCs (e.g. phospholipids remodeling or PUFA biosynthesis) could underlie qNSC aging. We focused on 5 lipid remodeling enzymes: MBOAT2, AGPAT3, and PLA2G4E (enzymes involved in phospholipid remodeling pathway) and ELOVL5 and FADS2 (enzymes in the PUFA biosynthesis pathway) (Fig. 4a). Using lentiviral-mediated CRISPR/Cas9, we knocked out the genes for each of these enzymes in primary cultures of qNSCs from young and old mice and performed untargeted lipidomic analysis (Fig. 4b). We verified that the knockout of these enzymes was efficient, with knockout qNSC cultures exhibiting frameshift editing in 55-70% of the alleles for each target gene (Extended Data Fig. 6a). Consistently, lipidomic analysis of the knockout qNSC cultures revealed the expected increase and decrease in substrate and product levels, respectively (Extended Data Fig. 6b, c, Supplementary Table 2).

Principal component analysis (PCA) on the lipidomic data showed that each gene knockout led to a distinct lipidomic profile in qNSCs (Fig. 4c). To assess the impact of the 5 lipid remodeling enzymes on the “youthfulness” of the lipidome, we leveraged the lipidomic signatures (based on membrane lipid side chain unsaturation and individual lipid features) we had generated above (see Fig. 3e). We generated a “lipid aging score” using these lipidomics signatures on lipidomics data for all 5 knockouts (see Methods). Interestingly, *Mboat2* knockout in old qNSCs led to a higher lipid aging score (“older”) compared to control, based on lipidomic aging signatures (combined features, Fig. 4d; side chain unsaturation features alone, Extended Data Fig. 6d). *Agpat3* knockout in old qNSCs also led to a higher score (“older”) compared to control, based on lipidomic aging signatures (combined features, Fig. 4d; individual lipid signatures alone, Extended Data Fig. 6e). Knockout of the other enzymes did not significantly affect lipidomic aging signatures. Thus, deficiency in the lysophospholipid acyltransferase MBOAT2 leads to changes in the membrane lipidomes that are associated with exacerbated aging in old qNSCs.

To functionally test the role of MBOAT2 in qNSC aging, we assessed the ability of qNSC to activate (i.e. transition from quiescence to proliferation), a key function of qNSCs that deteriorate with age^15, 46, 47, 88^. To this end, we knocked out *Mboat2* in qNSCs and assessed the proportion of qNSCs that activate in response to growth factors, using the proliferation marker Ki67 (Extended Data Fig. 8b). Old qNSCs were defective in their ability to activate compared to their young counterparts (Fig. 4e), consistent with previous observations^15, 46, 47, 88^. *Mboat2* knockout in old qNSCs led to a further impairment in qNSC activation ability (∼50% decrease, Fig. 4e). *Mboat2* knockout in qNSCs led to slight (not significant) decrease in plasma membrane order measured by Laurdan staining (Fig. 4f), though this assay may not be sensitive enough to detect subtle changes in membrane properties especially when other lipids are also remodeled. To test if *Mboat2* expression levels are themselves impacted by age, we re-analyzed our single-cell RNA-seq data from young and old SVZ neurogenic niches^14^. This analysis showed that *Mboat2* expression level decreased with age in qNSCs *in vivo* (Extended Data Fig. 6f). Together, these results suggest that deficiency in the phospholipid acyltransferase MBOAT2 with age could in part drive the decline in qNSC function during aging.

## Supplementation of plasma membrane lipids from young NSCs boosts old NSC function

Membrane lipids can be directly added to cells, without the need to manipulate specific genes. To assess the direct effects of plasma membrane lipids on old qNSC function, we supplemented old qNSCs with lipids extracted from plasma membrane of young qNSCs. We extracted lipids from GPMVs from young or old qNSC primary cultures (donor cells). We then supplemented these membrane lipids to a separate set of young or old qNSC cultures (recipient cells) (Fig. 5a). We verified successful uptake of plasma membrane lipids by recipient cells 72 hours after supplementation (Extended Data Fig. 7). While supplementation of plasma membrane lipids from old donor cells slightly reduced the activation of young recipient qNSCs (not significant), supplementation of plasma membrane lipids from young donor cells significantly boosted the ability of old qNSCs to activate (Fig. 5b).

**Figure 5.**
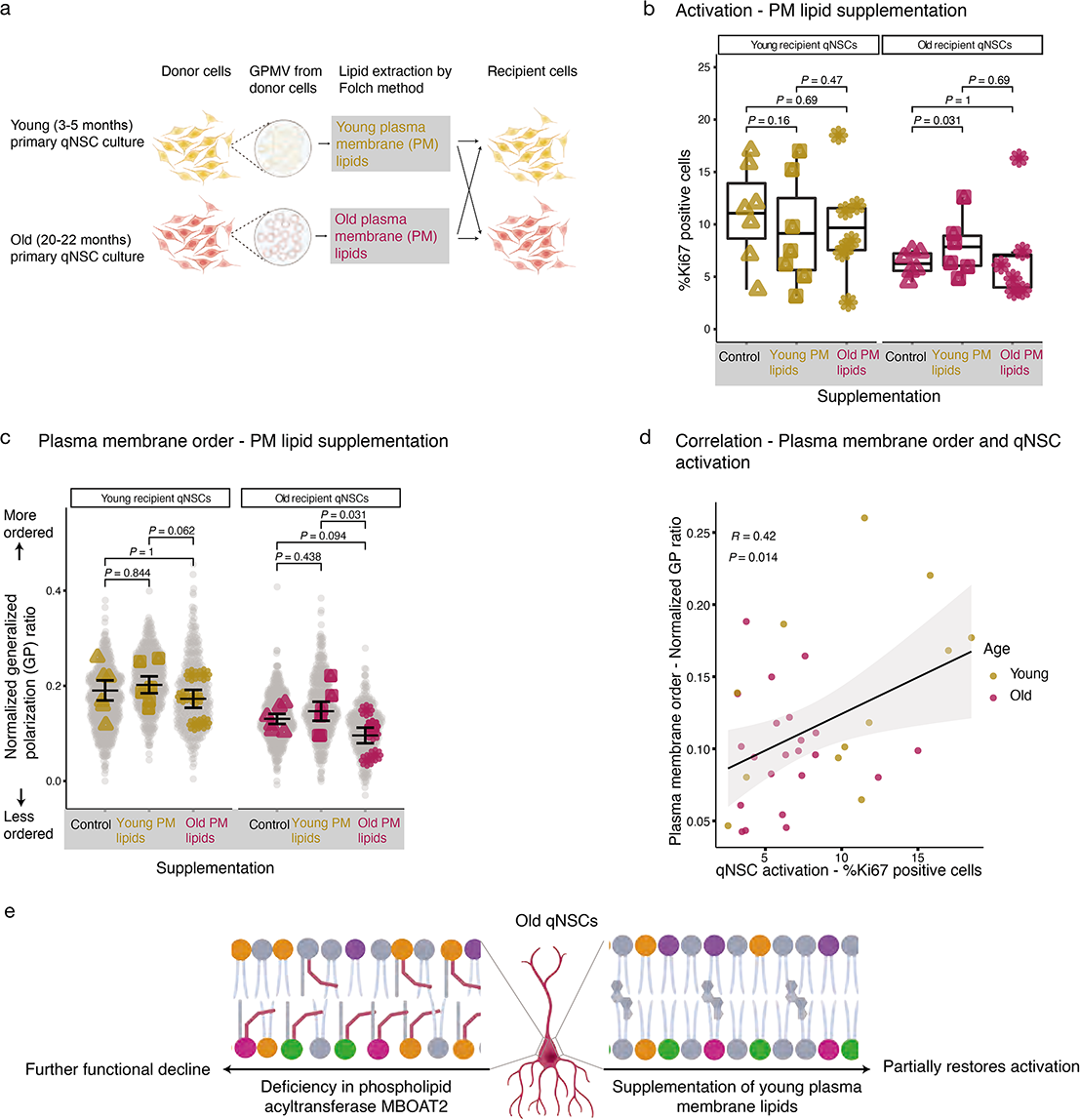
Supplementation of plasma membrane lipids from young cells boosts the activation ability of old quiescent neural stem cells. **a,** Overview of the experimental scheme: GPMVs were generated from young (3-5 months) or old (20-22 months) qNSC primary cultures (donor cells). Plasma membrane lipids from respective donor cells were extracted using a modified Folch method and then supplemented to a different set of young (3-5 months) or old (20-22 months) qNSC cultures (recipient cells) for assessment of activation and plasma membrane order. **b,** Activation (proliferation) ability of qNSCs with plasma membrane lipid supplementation. Primary culture of qNSCs from young (3-5 months) and old (20-22 months) mice were incubated with either vehicle control (methanol) or plasma membrane lipids from young (3-5 months) or old (20-22 months) donor qNSCs for 3 days. qNSCs were then switched to proliferative NSC media for 4 days. The percentage of young and old qNSCs that successfully activated was assessed by intracellular FACS analysis for Ki67, a proliferation marker. Box and whisker plot from 2 independent experiments, n=6-7 primary cultures. Horizontal bar: median. Whisker: Distance between the first/third quartile to the highest/lowest value, respectively, or 1.5 times the distance between the first/third quartile, whichever is smaller. Each dot represents a primary culture from an individual mouse. *P*-values from Wilcoxon signed-rank test (because of paired control and lipid supplementation). **c,** Plasma membrane order of young and old qNSCs upon plasma membrane lipid supplementation. Primary qNSC cultures from young (3-5 months) and old (20-22 months) mice were incubated with either vehicle control (methanol), plasma membrane lipids from young (3-5 months) or old (20-22 months) donor qNSCs for 3 days. Plasma membrane order was quantified by generalized polarization ratio (GP ratio) from Laurdan staining. SuperPlots showing data from individual cells (grey dots), as well as the mean of each primary culture (colored dots) +/-SEM. Results from two independent experiments, n=6 primary cultures. *P*-value from Wilcoxon signed-rank test (because of paired control and lipid supplementation). **d,** Correlation between plasma membrane order and activation in young and old qNSCs upon plasma membrane lipid supplementation. Pearson correlation was performed on generalized polarization ratio (GP ratio) from Laurdan staining, and percentage of qNSCs that activate (% Ki67 positive) from each individual young qNSC culture (golden dots) or old qNSC cultures (pink dots) with different supplementations. Linear regression line (black line) with 95% confidence interval (gray shade). **e,** Model for the functional impact of membrane lipids in quiescent neural stem cell aging.

In parallel, we determined the effects of plasma membrane lipid supplementation on plasma membrane order in young and old recipient cells (Fig. 5c). While the effects of supplemented lipids on plasma membrane order measured by Laurdan staining were small, supplementation of young membrane lipids led to a significant increase in plasma membrane order in old qNSCs (Fig. 5c). Overall, there was a positive correlation between recipient cells’ ability to activate and their plasma membrane order in both young and old qNSCs (Fig. 5d). Thus, supplementing lipids extracted from young plasma membranes boosts the function of old quiescent NSCs and increases membrane order in these cells. Together, these results suggest that lipid supplementation could be a strategy to counter NSC aging.

## Discussion

Our global lipidomic study on different mass spectrometry platforms shows that complex lipids strikingly change with age in old quiescent NSCs *in vitro* and *in vivo*. We find that many membrane lipids are remodeled with age and exhibit a shift in the side chain composition towards a higher PUFA content in quiescent NSCs. Our *in situ* lipidomic profiling in the SVZ neurogenic niche using DESI-MSI confirms age-related changes in membrane lipids of quiescent NSCs. By performing lipidomic analysis on isolated plasma membrane, we find that plasma membrane lipids largely contribute to these age-related lipidomic changes. Together, these comprehensive datasets allow us to generate robust “lipidomic aging signatures”^89^ that discriminate between young and old quiescent NSCs and can be leveraged to predict the impact of specific interventions. Prior lipidomics analyses have been conducted in cultured cells (e.g. oligodendrocytes, neurons, microglia, etc.)^30, 63, 73, 90–95^, but not in the context of organismal aging. In parallel, lipidomics studies have also been performed on whole organisms or organs (e.g. whole brains) in aging or Alzheimer’s disease^29, 31–34, 64, 89, 96–101^, but without cellular resolution. Thus, our lipidomics datasets represent the first systematic examination of the global lipidome of aging NSCs, and more generally of aging cells, *in vitro* and *in vivo*.

We observe a reduced plasma membrane order (more fluid) in quiescent NSCs. Membrane order has been shown to impact a broad range of biological processes, including immune signaling, virus infection, and signal transduction^24, 25, 102^. Reduced membrane order has also been observed in lymphocytes and neutrophils with age^103, 104^, whereas increased in membrane order was shown in hippocampal neurons with age^105^. These different changes in membrane properties with age may be directly linked to the function of the cell, its proliferative status^106^, and cell-cell contacts^107^. In old quiescent NSCs, a reduced membrane order may impair activation efficiency through disrupting the localization of transmembrane proteins important for proliferation. Indeed, plasma membrane lipids directly interact with a vast number of membrane proteins^24, 108, 109^, including transmembrane receptors, and aging may disproportionally affect membrane proteins^110–113^. Changes in plasma membrane properties may also affect other key aspects of quiescent NSC function, including endocytosis and phagocytosis.

We also identify interventions that affect plasma membrane lipids and find that they regulate the ability of quiescent NSCs to activate during aging (Fig. 5e). For example, deficiency in the phospholipid acyltransferase MBOAT2 further exacerbates aging defects in quiescent NSCs. Dysregulation of phospholipid acyltransferases has previously been linked to aberrant intestinal stem cell proliferation^114^, tumorigenesis^114, 115^ and liver diseases^116–118^. Our results now implicate a phospholipid acyltransferase in aging. Other enzymes, such as fatty acid elongases^119^, may also contribute to aging defects. Interestingly, we also show that supplementing young plasma membrane lipids boosts activation in old qNSCs and that plasma membrane order correlates with the ability of qNSCs to activate. Thus, restoring specific lipids to a young lipid profile – either genetically or by affecting lipids themselves – could be an effective strategy to slow cellular aging and counter age-related diseases, including neurodegenerative diseases.

## Acknowledgments

We thank Katharina Papsdorf, Lucy Xu, Jingxun Chen, Param P. Singh, Rahul Nagvekar, Jason W. Miklas and all Brunet lab members for their input on the project and for providing feedback on the manuscript. We thank Itay Budin and Bianxiao Cui for helpful discussion on membrane lipid assays and Jonathan Z. Long for functional lipid experiment designs. We also thank members of the Cui lab and Snyder lab for advice on the project. This work was supported by NIH P01AG036695 (A.B.), a generous gift from Timothy and Michele Barakett, AFOSR FA9550-21-1-0170 (R.N.Z), and a postdoctoral fellowship from American Federation for Aging Research (X.Z.). Schematic diagrams were created with BioRender.com.

## Author contributions

X.Z. designed the project with help from A.B.. X.Z. performed all experiments and computational analyses unless otherwise indicated. X.Y. performed DESI-MSI lipidomic profiling and structural validation under the supervision of R.N.Z.. K.C. helped optimize and run lipidomic studies on LC-MS/MS, provided input on data validation and analysis under the supervision of M.P.S., and helped with independent code-checking. F.V. performed cell type-specific deconvolution on DESI-MSI data, and provided input on lipidomic aging signatures under the supervision of P.K.. M.E. helped run LC-MS/MS lipidomic samples and data analysis under the supervision of M.P.S.. C.M.K. helped with CRISPR/Cas9 knockout in NSCs and data analysis on Laurdan staining. S.D.G. performed western blot experiments. C.J.S. helped optimize CRISPR/Cas9 knockout and activation assay in NSCs. M.C. helped with *in vivo* sorted lipidomic samples, data analysis, and independent code checking under the supervision of M.P.S.. G.M.T. helped optimizing and running LC-MS/MS lipidomic samples under the supervision of M.P.S.. A.M. performed image analysis for DESI-MSI dataset and helped with independent code-checking. D.H. helped run and analyze data from Lipidyzer platform under the supervision of M.P.S.. M.P.S. and R.N.Z. gave intellectual input on the study. X.Z. wrote the manuscript with A.B.. All authors provided intellectual input and commented on the manuscript.

## Competing interests

The authors declare no competing interests.

## Data availability

Raw lipidomics data were uploaded to Metabolomicsworkbench.org under Study ID ST002257 (DatatrackID:3404), Study ID ST002258 (DatatrackID:3406), Study ID ST002259 (DatatrackID:3407) and Study ID ST002260 (DatatrackID:3408). Processed lipidomics data are included as Supplementary Tables 1-6.

## Code availability

The code used to analyze lipidomics data in the current study is available in the Github repository for this paper (https://github.com/xiaoaizhao/Neural-stem-cell-NSC-aging-lipidomics).

## Methods

### Laboratory animals

All mice used in this study were male C57BL/6JN mice (hereafter C57BL/6) obtained from the NIA Aged Rodent colony. Mice were habituated for more than two weeks at Stanford before use. At Stanford, all mice were housed in the Comparative Medicine, and their care was monitored by the Veterinary Service Center at Stanford University under IACUC protocol 8661.

### Primary cultures of quiescent and activated NSCs

For primary cultures of mouse NSCs, NSCs were isolated from male C57BL/6 mice as previously described^1–3^. Briefly, subventricular zones (SVZs) from each brain were microdissected and finely minced. Tissue suspension was then digested for 35min at 37°C with gentle agitation in HBSS media (Corning, 21-021-CV) containing 2U/ml Papain (Worthington LS003124), 1U/ml Dispase II (STEMCELL Technologies, 07913), and 0.1mg/ml DNase I (Sigma, DN25-100mg), and mechanically dissociated. Isolated cells were expanded as neurospheres in culture in “Proliferative NSC media” (NeuroBasal-A medium (Gibco, 10888-022) with penicillin-streptomycin-glutamine diluted 1X (Gibco, 10378-016), 2% B27 minus vitamin A (Gibco, 12587-010), 20ng/ml bFGF (Peprotech, 100-18B) and 20ng/ml EGF (Peprotech, AF-100-15)) at 37°C in 5% CO_2_ and 20% O_2_ at 95% humidity.

To generate parallel cultures of quiescent and activated NSCs (qNSCs and aNSCs, respectively), we used a previously described protocol^4^. Specifically, 1×10^6^ NSCs (proliferating, passage 3 to passage 5) were plated in each well of a 6-well plate (80-90% density). To generate primary cultures of quiescent NSCs (qNSCs), tissue culture plates were pre-treated with PBS (Corning, 21-040-CV) containing 50ng/ml Poly-D-Lysine (Sigma-Aldrich, P6407) for 2 hours in 37°C tissue culture incubator, and then washed 3 times with PBS prior to plating cells. NSCs were then cultured in “Quiescence NSC media” (NeuroBasal-A (Gibco, 10888-022), penicillin-streptomycin-glutamine 1X (Gibco, 10378-016), 2% B27 minus vitamin A (Gibco, 12587-010), 20ng/ml bFGF (Peprotech, 100-18B) and 50ng/ml BMP4 (Biolegend, 595302). For lipidomics analysis, generation of giant plasma membrane vesicles (GPMVs) and membrane order assay by Laurdan staining, qNSCs were incubated in this quiescence media for 7 days before the experiment. For CRISPR/Cas9 knockout, qNSCs were incubated in quiescent media for 4 days before 2 additional days of lentiviral transduction in the same media. For all experiments on qNSCs, quiescence media was replaced every 2 days.

To generate primary cultures of aNSCs, 700,000 NSCs (proliferating, passage 3 to passage 5) were plated in each well of a 6-well plate pre-treated with poly-D-lysine as described above. Cells were incubated in “Proliferative NSC media” (see above) for 2 days before harvesting for lipidomic analysis. The difference in the number of plated cells between qNSCs and aNSCs is to ensure similar cell density in both quiescence and activated conditions at the time of harvesting.

### Primary NSC culture sample preparation for lipidomics

For lipidomics, aNSCs and qNSCs were washed twice with PBS before incubating in “Proliferative NSC media” minus B27 supplement and “Quiescent NSC media” minus B27 supplement, respectively. Cells were incubated for 3 hours in these media at 37°C incubator with 5% CO_2_ and 20% oxygen at 95% humidity to remove exogenous lipids contained in B27 supplement. At the end of the incubation period, cells were washed once with PBS 1X (Corning, 21-040-CV) and scraped into 500μl of ice-cold PBS using cell lifter (Fisher Scientific 07-200-364). The cell suspension was collected in 2ml amber glass vials (Thermo Scientific, 03-FISVA) sealed with polyethylene cap with PTFE/silicone septum (Waters, 186000274). All samples were immediately snap-frozen in liquid nitrogen and stored at −80°C.

### Lipid extraction for lipidomics analysis by liquid chromatography followed by tandem mass spectrometry (LC-MS/MS)

Lipids were extracted from cell suspension after thawing on ice using a modified Folch method^5^. All chemical reagents used were LC-MS grade unless indicated otherwise. Specifically, 300μl of cold 100% methanol (Fisher Scientific, A456-500) containing deuterated lipid standards listed below was added to the cell suspension. Deuterated triacylglycerol TG(17:0-17:1-17:0(d5)) (Avanti Polar Lipids, 860903) 0.1μg was used for normalization in the untargeted LC-MS/MS analysis for lipidomics on activated and quiescent primary NSC culture (**primary NSC culture experiment #1**). A mixture containing 54 deuterated standards (SCIEX, 5040156, LPISTDKIT-101) was used for the **targeted Lipidyzer analysis** (20μl/sample). A mixture containing 13 deuterated standards (EquiSPLASH® mix, Avanti Polar Lipids, 330731) and deuterated oleic acid (Cayman Chemical, 9000432) was used for the untargeted LC-MS/MS analysis (1μl/sample) on quiescent NSC cultures with CRISPR/Cas9 knockout (**primary NSC culture experiment #2**). Homogenates were sonicated three times for 30s each time at room temperature in a water bath sonicator (VWR, 97043-960). Samples were rested on ice for 30s between each cycle. Following this step, 600μl of cold chloroform (Acros Organics, AC610281000, stored at −20°C) was added to the homogenates. Samples were then subjected to vigorous vortex at 4°C for 30min. Biphasic separation was achieved after centrifugation at 3000rpm for 10min at 4°C. The lower organic phase containing the lipids was collected and dried down under a nitrogen stream using a TurboVap Classic LV (Biotage) at a flow rate of 0.5l/min for 15min until no visible solution remains, and with dried lipid film formed at the bottom. Dried lipids were then resolubilized in 200μl 100% methanol at room temperature before moving to −20°C for storage. On the day of analysis, for untargeted LC-MS/MS, half of each sample’s lipid extract was dried down under a nitrogen stream and resolubilized in 200μl of methanol:toluene (90:10, vol:vol) for analysis on complex lipids, and the other half of the lipid extract was resolubilized in 100μl of 5% acetonitrile for free fatty acid analysis. For targeted assay on the Lipidyzer platform, samples were solubilized in 300μl of 10mM ammonium acetate in methanol:toluene (90:10, vol:vol) before analysis.

### Untargeted lipidomics by LC-MS/MS

Lipid extracts were analyzed in a randomized order using an Ultimate 3000 RSLC system coupled with a Q Exactive mass spectrometer (Thermo Scientific) as previously described^6^. Each sample was run twice in positive and negative ionization modes. Lipids were separated using an Accucore C18 column 2.1 x 150mm, 2.6μm (Thermo Scientific, 17126-152130) and mobile phase solvents consisted of 10mM ammonium acetate and 0.1% formic acid in 60/40 acetonitrile/water (A) and 10mM ammonium acetate and 0.1% formic acid in 90/10 isopropanol/acetonitrile (B). The gradient profile used was 30% B for 3min, 30-43% in 2min, 43-55% B in 0.1min, 55-65% in 10min, 65-85% B in 6min, 85-100% B in 2min and 100% B for 5min. Lipids were eluted from the column at 0.4ml/min, the oven temperature was set at 45°C, and the injection volume was 5μl. Autosampler temperature was set at 20°C to prevent lipid aggregation. The Q Exactive was equipped with a HESI-II probe and operated in data-dependent acquisition mode for whole cell samples with CRISPR/Cas9 knockouts. To increase sensitivity in untreated whole cell and giant plasma membrane vesicles (GPMVs) samples (see below), samples were run in full MS mode and MS/MS spectra were acquired on pooled samples. To maximize the number of identified lipids, the 100 most abundant peaks found in blanks were excluded from MS/MS events. External calibration was performed using an infusion of Pierce LTQ Velos ESI Positive Ion Calibration Solution or Pierce ESI Negative Ion Calibration Solution. Data quality was ensured by 1) injecting 6 pooled samples to equilibrate the LC-MS system prior to run the sequence, 2) checking mass accuracy, retention time, and peak shape of internal standards in each sample.

### Free fatty acid analysis by LC-MS

Free fatty acids were analyzed in a randomized order using an Ultimate 3000 RSLC system coupled with a Q Exactive mass spectrometer (Thermo Scientific). The Q Exactive was operated in data-dependent acquisition mode in negative ionization mode. Free fatty acids were separated on a Zorbax StableBond aq column (dimension: 2.1mm x 50mm, particle size: 1.7µm) (Agilent technologies, 827700–914) at 0.6ml/min using a mobile phase solvent consisted of 0.06% acetic acid in water (A) and 0.06% acetic acid in acetonitrile (B). The gradient was set with 1-90%B in 9min, the oven temperature was set at 60°C, the autosampler temperature was set at 4°C, and the injection volume was 15μl. Single-point internal standard calibration with deuterated oleic acid (Cayman Chemical, 9000432) was used to estimate absolute concentrations for all free fatty acids.

### Nomenclature

Lipid nomenclature used here follows the standardized guidelines^7^. The number of C=C bonds is indicated by the number after the carbon number in the acyl chain and is separated by “:”. The separator “_” is used between the two acyl chains if sn-position of the fatty acids is not known. The separator “/” is used when sn-position of fatty acids is proven (e.g., sn-1/ sn-2 or sn-2/ sn-3 for glycerophospholipids).

### LC-MS/MS data processing

Data from each mode were independently analyzed using Progenesis QI software (v2.3, Nonlinear Dynamics). Metabolic features from blanks and that did not show sufficient linearity upon dilution in QC samples (r < 0.6) were discarded. Only metabolic features present in >2/3 of the samples were kept for further analysis. Lipids were identified using LipidSearch (v4.3, Thermo Scientific) by matching the precursor ion mass to a database and the experimental MS/MS spectra to a spectral library containing theoretical fragmentation spectra. The most abundant ion adduct was selected for each lipid class for downstream analysis and quantification. Specifically, in positive mode, [M (molecular ion) +H]+ for Lysophosphatidylcholine (LPC), Phosphatidylcholine (PC), Phosphatidylethanolamine (PE), Sphingomyelin (SM), Acylcarnitine (AcCa) and Coenzyme (Co), [M+NH_4_]+ for Cholesterol ester (ChE), Monoacylglycerol (MG), Diacylglycerol (DG) and Triacylglycerol (TG), [M+H-H_2_O]+ for Ceramide (Cer) and Cholesterol. In negative mode, [M-H]-for Lysophosphatidylethanolamine (LPE), Phosphatidylinositol (PI), Phosphatidylserine (PS), Phosphatidylglycerol (PG), Cardiolipin (CL), Lysophosphatidylinositol (LPI) and Lysophosphatidylserine (LPS). To reduce the risk of misidentification, MS/MS spectra from lipids of interest were manually investigated to validate the assignments. The manual validation consisted in verifying that: 1) both positive and negative mode MS/MS spectra matched the expected fragments, 2) the main lipid adduct forms detected in positive and negative modes were in agreement with the lipid class identified, 3) the retention time was compatible with the lipid class identified, and 4) the peak shape was acceptable. The fragmentation pattern of each lipid class detected was experimentally validated using lipid internal standards.

In primary NSC culture experiment #1, internal standard (TG(17:0-17:1-17:0(d5)), see above) - normalized signal intensity was obtained for all detected lipids. Subsequently, median lipid intensity of each sample was used to normalize for variation in starting material before performing downstream analyses. In primary NSC culture experiment #2, the inclusion of EquiSPLASH® deuterated lipid standard mix (see above) allowed us to obtain quantitative molar concentration for identified lipids that belong to the 13 lipid classes (PC, LPC, PE, LPE, PG, PI, PS, TG, DG, MG, ChE, Cer, SM). For those lipids, single-point internal standard calibrations were performed to estimate absolute concentrations for each lipid. A number of less abundant lipid classes of Cholesterol (Chol), Cardiolipin (CL), Acyl carnitine (AcCa), Co-enzyme (Co), Sphingosine phosphate (SPHP) and Zymosterol ester (ZyE) were also detected in primary NSC culture experiment #2. As these lipids do not have internal standard for their respective lipid classes, normalized signal intensity was obtained instead of molar concentration. Normalized intensity for each lipid was calculated using the median lipid molar concentration of quantified lipids of each sample to normalize for variation in starting material. Importantly, we ensured linearity within the range of detected endogenous lipids by using serial dilutions of deuterated standards spanning 4 orders of magnitude. Subsequently, median lipid molar concentration of each sample was used to normalize for variation in starting material.

### Targeted lipidomics using the Lipidyzer platform

Lipid extracts were analyzed using the Lipidyzer platform that comprises a 5500 QTRAP System equipped with a SelexION differential mobility spectrometry (DMS) interface (SCIEX) and a high flow LC-30AD solvent delivery unit (Shimazdu) as previously described^6^. Briefly, lipid molecular species were identified and quantified using multiple reaction monitoring (MRM) and positive/negative ionization switching. Two acquisition methods were employed covering 10 lipid classes; method 1 had SelexION voltages turned on while method 2 had SelexION voltages turned off. Lipidyzer data were reported by the Lipidomics Workflow Manager (LWM) software which calculates concentrations for each detected lipid as average intensity of the analyte MRM/average intensity of the most structurally similar internal standard (IS) MRM multiplied by its concentration. Lipid abundance was reported as concentration in nmol/ml. Similar to untargeted LC-MS/MS data, median lipid molar concentration of each sample was used to normalize for variation in starting material before performing downstream analyses. Data quality was ensured by: 1) tuning the DMS compensation voltages using a set of lipid standards (SCIEX, 5040141) after each cleaning. Tuning was also performed when the system has been idling for more than 24 hours, or after 3 consecutive days of running. 2) performing a quick system suitability test (QSST) (SCIEX, 50407) and 3) triplicate injection of lipids extracted from a reference plasma sample (SCIEX, 4386703) before data acquisition to ensure acceptable limit of detection for each lipid class.

### Heatmap visualization on lipids with age-related changes in primary quiescent NSC culture

Differential lipids with age in quiescent NSC culture were obtained using a false discovery rate (FDR) < 0.05. Heatmap was generated from differential lipids (95 lipids total) with Pearson correlation using the R package “ComplexHeatmap” v2.6.2.

### Double bond composition analysis

For each individual lipid class, we grouped lipids based on the presence of double bond 0-6 in any of their identified side chains. Normalized lipid intensity was used in the following datasets/subset of data – all lipids in primary NSC culture experiment #1, lipid classes detected but without corresponding deuterated internal standards in primary NSC culture experiment #2, *in vivo* sorted qNSC and giant plasma membrane vesicles (GPMVs). Normalized lipid concentration was used in the following datasets/subset of data – lipid classes with corresponding deuterated internal standards in primary NSC culture experiment #2, *in vivo* sorted qNSC and giant plasma membrane vesicles (GPMVs). To calculate the composition of each double bond number within a given class, the aggregated lipid intensity/concentration is divided by the total intensity/concentration of the same class. Only lipid classes that have 10 or more detected lipids were included in this analysis.

### *In vivo* isolated NSC lipidomics

To isolate quiescent neural stem cells (qNSCs) directly from mouse brain, we used fluorescence-activated cell sorting (FACS)^2, 3^. Briefly, the subventricular zones (SVZs) of young (3-5 month-old) and old (20-22 month-old) C57BL/6 male mice were micro-dissected following a previously published protocol^8^. Single cell suspension was obtained and then stained with 1:50 CD31 (BioLegend, 102405), 1:50 CD45 (BioLegend, 103108), 1:50 CD24 (BioLegend, 101805), 1:300 Prominin-1-biotin (Thermo Scientific, 13-1331-82) and 1:300 EGF-Alexa 647 (Molecular probes, E35351) for 30min on ice. Cells were then stained with 1:1000 Streptavidin-PEcy7 (eBioscience, 25-4317) for 30min on ice. Quiescent NSCs (CD31^-^CD45^-^CD24^-^Prominin-1(CD133)^+^EGFR^-^) were sorted. All FACS sorting was performed at the Stanford FACS facility on a BD Aria sorter, using a 100μm nozzle at a pressure of 13 psi. We pooled 5 young mice or 5 old mice to obtain ∼7,500-9,000 qNSCs for each biological replicate. qNSCs were sorted into 500μl of ice-cold PBS, and the cell suspension was immediately snap-frozen in liquid nitrogen and then transferred to −80°C for storage. As previously described in untargeted lipidomics in primary NSC culture #2, a mixture containing 13 deuterated standards (EquiSPLASH® mix, Avanti Polar Lipids, 330731) was added to each sample (0.5μl/sample), together with 0.1μl of Cholesterol (d7) (Avanti Polar Lipids, 700041). Single-point internal standard calibrations were used to estimate absolute concentration for identified lipids using one internal standard for each lipid class. Subsequently, median lipid molar concentration of each sample was used to normalize for variation in starting material.

### Principal Component Analysis (PCA) on lipidomics datasets

All identified lipids were used to visualize global lipidome of each individual samples. Principal Component Analysis (PCA) was conducted on normalized lipid intensity (primary NSC culture experiment #1). For primary NSC culture experiment #2 and *in vivo* sorted qNSC dataset, normalized lipid molar concentration was used for lipid classes with deuterated internal standard, while normalized lipid intensity was used for all other lipid classes (see section on LC-MS/MS data processing). For *in situ* qNSC lipidomic profiling by DESI-MSI, normalized metabolite intensity was used for this analysis. PCA was generated using function “prcomp()” with R version 4.0.2. To visualize PCA plots comparing young and old samples of the same cell type, we selected the top two components ranked by their area under the curve of a receiver operator curve (AUC) in differentiating age differences. We computed a decision boundary in the plot by applying a logistic regression on the two chosen components.

### Effect size calculation

Hedges’ *g* statistics was used to evaluate the standardized mean difference between two groups for a given measurement^9, 10^. Specifically,

*g* = correction factor * ((Mean_1_-Mean_2_)/SD_weighted_pooled, where SD_weighted_pooled = sqrt(((n_1_-1)*SD ^2^ + (n_2_-1)*SD ^2^)/(n_1_+n_2_-2))

A correction factor was applied to experiment with small sample size (<50 samples). correction factor = 1-3/(4*(n_1_+n_2_)-9)

Standard error = sqrt((n_1_ + n_2_)/(n_1_ * n_2_) + 0.5 * *g*^2 /(n_1_ + n_2_-3.94))

### SVZ sample preparation for Desorption Electrospray Ionization Mass Spectrometry Imaging (DESI-MSI)

SVZ neurogenic niches were micro-dissected from 7 young (3-5 month-old) and 7 old (20-22 month-old) C57BL/6 male mice brains. The tissue was oriented on a cryomold (Fisher scientific, NC9511236) such that the lateral wall of the SVZ faces down to flatten the tissue, before immediately snap freezing the sample on dry ice. For cutting, SVZs were not embedded in optimal cutting temperature (OCT) polymeric compound to avoid background compound peaks from residual OCT during tissue sectioning. Instead, a minimal amount of OCT was applied to attach the caudoputamen, which is adjacent to the lateral wall of the SVZ, to the mounting plate before sectioning. Unfixed SVZs were cut into 16μm sections parallel to the lateral wall of the SVZ at −20°C on a Leica CM3050S cryostat. Sections were thaw-mounted onto slides (Fisher scientific, 12-550-15) without fixation, before transferring to −80°C for storage. Tissue slides were dried for 20min at room temperature using a vacuum desiccator (Thermo scientific, 5311-0250) before DESI-MSI experiments.

### DESI-MSI

The DESI-MSI procedure was previously described^11^. Briefly, a custom-built DESI imaging stage was coupled to a hybrid LTQ-Orbitrap mass spectrometer (Thermo Fisher Scientific) for DESI-MSI. The mass spectra were acquired in the negative ion mode from *m/z* 50−1000 using the Orbitrap mass analyzer with the resolving power set at 60,000. The spray voltage was set to −3.5kV, and the capillary voltage was set to −65V. The tube lens voltage was set to −120V. Ion injection time was 100ms, and one microscan was performed. The solvent system used is *N*,*N*-dimethylformamide (DMF)/ acetonitrile (ACN) = 1:1 (vol/vol) and provided at a flow rate of 0.8μl/min. Assisted by a nebulizing gas (N_2_) at a pressure of 150psi, molecules of interest from tissues are desorbed and ionized. DESI spray tip-to-surface distance was 2mm; spray incident angle was 56°, and spray-to-inlet distance was 6.5μm. Step size in the moving stage was set to 200μm, and automatic gain control of mass spectrometer was switched off. These parameters were empirically found to yield the optimal MS signal from brain tissues. Tandem mass spectra were obtained using collision-induced dissociation from the ion trap mass analyzer to validate lipid structures based on their characteristic fragment ions. All experiments were carried out under identical experimental conditions to allow comparison between measurements.

### Immunofluorescence staining and imaging for DESI-MSI

After each DESI scan, tissue sections were fixed with 4% paraformaldehyde (Electron Microscopy Science, 15714) diluted in phosphate-buffered saline for 20min at −20°C. Antibodies against Glial Fibrillary Acidic Protein (GFAP) (Abcam, ab53554) were used at 1:1000 dilution, antibodies against Ki-67 (eBioscience, SolA15) were used at 1:500 dilution at 4°C overnight. Donkey anti-goat antibodies conjugated with Alexa Fluor 647 (Thermo Fisher Scientific, A-21447), and donkey anti-rat antibodies with Alexa Fluor 488 (Thermo Fisher Scientific, A-21208) was used at 1:250 dilution at room temperature for 1.5h for a secondary staining. All sections were mounted with 50μl of ProLong Gold Antifade mounting media containing a fluorescent nucleic acid dye 4′,6-diamidino-2-phenylindole (DAPI) (Thermo Fisher Scientific, P36931) before imaging. Confocal microscopy was carried out using a Nikon Eclipse Ti confocal microscope equipped with a Zyla sCMOS camera (Andor) and NIS-Elements software (AR 4.30.02, 64-bit) using a 10X objective. A tiled image covering the entire area of SVZ was taken for each mouse.

### Image quantification

Image quantification was performed using a Julia language script. Nuclei-detection was performed for immunofluorescence-stained regions of tissue using DAPI as a nuclear marker and blob detection algorithms provided by the Julia package Images.jl. After nuclei detection, cells were classified as either GFAP-positive and Ki67-positive (activated NSCs), GFAP-positive and Ki67-negative (quiescent NSCs/astrocytes), GFAP-negative and Ki67-negative (likely neurons), based upon the staining in the region immediately surrounding the detected nuclei. To threshold between these binary designations we used Otsu’s method^12^ on the summed staining intensity for a given channel by cell. After each cell was classified and binned into a category based upon the calculated threshold, each region’s proportion of cell types was calculated and presented as a percentage of total cells in that region for downstream analysis.

### Cell type-specific deconvolution of mass spectra

Peak intensity of each individual DESI scan was normalized by dividing each peak by the intensity value of an exogenous standard (4-methoxybenzoic acid) introduced by the solvent system. We then aggregated all normalized peak intensities by rounding their m/z value by its first significant digit to reduce data sparsity prior to deconvolution. We estimated cell-type specific intensity values using the csSAM package (version 1.2.4)^13^ by pairing normalized DESI data with cellular proportions quantified by immunofluorescent staining on the same sample, as described in the section above. Specifically, we grouped all scans obtained from the same animal to estimate mean and standard error across all peaks for each mouse. We then used cell-type specific mean intensities for statistical comparisons and downstream analyses between mice. Fold change differences between old and young mice were estimated by computing Hedges’ *g* effect size (described above) using age groups as class labels.

### Evaluation of cell type-specific lipidomic deconvolution accuracy

To validate the accuracy of cell type-specific deconvolution by csSAM, we generated a reconstituted *in silico* mixture lipidomic dataset. To best mimic the 4-way deconvolution applied to our DESI-MSI data, we used lipidomic data from 4 different cell types - quiescent and activated NSC from primary NSC culture experiment #1 (see above), and untargeted lipidomic data from cultures of two additional brain cell types – neurons and oligodendrocytes^14^. We then reconstituted mixture lipidomic data to simulate our DESI-MSI data samples. Specifically, we grouped one spectra from each cell type to generate an unique *in silico* lipidomic profile representing an individual mouse. We then created 10 mixtures for each mouse by combining the spectra of 4 cell types using a different vector of cell-type proportion for each mixture. We performed deconvolution on these reconstituted mixture lipidomic data, and obtained estimated mean levels of cell-type specific metabolite intensity for each individual mouse. Finally, we compared mean estimated metabolite intensity after deconvolution to the measured metabolite intensity prior to mixing by Pearson correlation.

As a negative control, we also performed deconvolution using the same set of reconstituted *in silico* mixture but with mismatched cell type proportions instead. Specifically, we performed deconvolution with randomly generated cell type compositions from 100 permutation experiments. We then analyzed the distribution of the Pearson correlation coefficient between the measured and estimated lipid intensity for each cell type of each individual mouse after deconvolution.

### Sparse partial least squares - discriminant analysis (sPLS-DA) and feature selection

After cell type-specific deconvolution, we identify metabolic signatures associated with either cell type or age group using sparse partial least squares - discriminant analysis (sPLS-DA), a supervised multivariate statistical method used to identify features that maximize the variance between dependent variables and class labels^15^. We performed sPLS-DA using the mixOmics package (version 6.14.1)^16^, specifying two components for our analysis to facilitate result interpretation and visualization. Briefly, for each component, we iteratively performed sPLS-DA specifying a number of features ranging from a minimum of 1 and a maximum of 2000. We then performed an elbow analysis to identify the smallest number of features that is able to provide the correct classification. Specifically, we selected features corresponding to an area under the curve of a receiver operator curve (AUC) that is the closest to the maximum AUC within ε = 0.01. We repeated this step for each component and then combined features from both components to obtain metabolites or lipids with age or cell type-specific changes.

### Visualization and analysis of age or cell type-specific metabolites in DESI-MSI

To identify age-related changes across cell types, we used the normalized peak intensity of all age-specific features identified by sPLS-DA (see above). Heatmap was then generated using the standard score (Z score) of each metabolite with Pearson correlation using the R package “ComplexHeatmap” v2.6.2. We assessed the degree of age/cell type interactions across all metabolites by two-way ANOVA followed by correction of multiple hypothesis testing. Significant interaction was identified using FDR < 0.2 due to the large number of metabolite features, for cell type-specific response with age.

### Giant plasma membrane vesicle (GPMV) generation and lipidomics

Giant plasma membrane vesicles (GPMVs) were generated from primary qNSC cultures from individual mice following a previously described protocol^17, 18^. Specifically, 13×10^6^ qNSCs (passage 5 to passage 6) were plated in a 15cm plate and quiescence was induced for 7 days (see above). Cells were washed twice with “GPMV vesiculation buffer” (150mM NaCl, 2mM CaCl_2_, and 20mM HEPES (Sigma-Aldrich, 54457) in water, pH7.4) at 37°C. GPMV generation was then induced by adding 2mM of *N*-ethylmaleimide (Sigma-Aldrich, 04259) for 1.5h in 37°C incubator with 5% CO_2_ and 20% O_2_, with gentle agitation every 30min. At the end of the incubation, the cell supernatants (containing GPMVs) were first subjected to a low speed centrifugation at 200g for 5min at room temperature to remove cellular debris. Supernatants were then carefully collected, followed by ultracentrifugation at 30,000g for 30min at 4°C using a Beckman Coulter, Optima XE-90 ultracentrifuge. After removing supernatant, pelleted GPMVs with ∼500μl of residual buffer were snap frozen in liquid nitrogen and transferred to −80°C for storage. As previously described in untargeted lipidomics for *in vivo* sorted qNSCs, a mixture containing 13 deuterated standards (EquiSPLASH® mix, Avanti Polar Lipids, 330731) was added to each sample (0.5μl/sample), together with 0.1μl of Cholesterol (d7) (Avanti Polar Lipids, 700041) for quantification and normalization. Similarly, single-point internal standard calibrations were used to estimate absolute concentrations for identified lipids using one internal standard for each lipid class. Subsequently, median lipid molar concentration of each sample was used to correct for variation in starting material.

### Western blot analysis

For whole cell lysate, 13×10^6^ qNSCs (passage 5 to passage 6) were plated in a 15cm plate and quiescence was induced for 7 days (see above). Cells were then lysed directly in the culture plates using 3ml of ice-cold 1X RIPA buffer diluted from 5X RIPA buffer (Thermo Scientific, J62524AD) with ultrapure water (Invitrogen, 10977-015) and supplemented with 1X protease inhibitor cocktail (Roche, 04693159001). Cell lysates were then scraped and transferred to Eppendorf tubes. GPMV were generated from 13×10^6^ qNSCs (passage 5 to passage 6) as described above. GPMV protein lysates were obtained by lysing GPMV pellet with 50μl of 1X RIPA buffer with 1X protease inhibitor cocktail. Following the addition of Laemmli sample buffer (Alfa, Aesar, J61337), all lysates were resolved on NuPAGE 4-12% gradient Bis-Tris gel (Invitrogen, NP0321BOX), transferred onto nitrocellulose membranes (BioRad, 162-0115) and immunoblotted. For each sample, equal amounts of total protein was loaded based on protein quantification using Ponceau S staining (see below). Primary antibodies used were: 1:25,000 Beta-actin (Abcam, Ab6276), 1:5000 Calnexin (Abcam, Ab22595) and 1:5000 Cox4 (Abcam, Ab16056). Membranes were then incubated with secondary antibody IRDye 680RD donkey anti-mouse IgG (LI-COR, 926-68072), IRDye 800CW goat anti-rabbit IgG (LI-COR, 926-32211), or IRDye 680RD goat anti-rat IgG (LI-COR, 926-68076). All secondary antibodies were used at 1:5000 dilution. Blots were imaged on Odyssey XF Imager (LI-COR).

### Ponceau S staining and total protein quantification

Following protein gel transfer onto nitrocellulose membranes, the membranes were incubated in Ponceau S solution (Sigma-Aldrich, P7170-1L) for 5min at room temperature and then washed 3 times (1min each time) with Milli-Q water. The membrane was immediately scanned on an Epson perfection V300 photo scanner (model #J232B) for densitometry quantification. Using ImageJ (version 2.1.0/1.53c), the images were split into red, blue, and green channels. Equal sized rectangles were drawn around each lane, and the mean integrated density from each lane of each channel was measured. The sum of the mean integrated density from all three channels was compared to verify the same amount of protein was loaded into each lane.

### Lipid ontology enrichment analysis

We performed lipid ontology (LION) enrichment analysis by inputting lipid lists into the LION/web online portal (http://www.lipidontology.com/), and then selected all terms under “cellular component”. A local statistic is computed for each lipid by LION using one-tailed Welch two-sample *t*-tests. Enrichment score for each ontology term is then generated by LION using two-tailed Kolmogorov–Smirnov tests^19^.

### Lipidomic aging signature

To identify the most robust and reproducible lipidomic changes with age *in vitro* and *in vivo*, we performed a meta-analysis across the following datasets - primary NSC culture experiment #1, primary NSC culture experiment #2, *in vivo* sorted qNSCs and GPMV lipidomics. For input, we calculated the age-related effect size (see above) on double bond composition for each lipid class obtained from our double bond composition analysis (see above) on all datasets. Meta-analysis was done by R package “rmeta” v3.0 with random effects model. Lipidomic aging signature was then selected using the following criteria: 1) overall *P* value <0.05, 2) same directional change detected in at least 3 out of 4 datasets, 3) homogeneity between datasets (*P* value heterogeneity >0.05), 4) summary effect size >0 (indicates increase with age).

### Laurdan staining and quantification in primary NSC culture

Membrane order was assessed using the polarity-sensitive membrane probe Laurdan (6-Dodecanoyl-2-Dimethylaminonaphthalene) (Thermo Fisher Scientific, D250)^20, 21^. Specifically, 90,000 qNSCs from primary culture of young and old mice were plated in each well of 8-well chamber coverslip (Ibidi, 80826). After 7 days of induced quiescence, cells were stained with Laurdan at a final concentration of 30μg/ml in culture media for 30min in 37°C tissue culture incubator. Cells were then washed once with media. DNA stain DRAQ5^TM^ (Thermo Fisher Scientific, 62251) was added at a final concentration of 5μM in media, immediately before imaging. Live cell imaging was carried out in 37°C imaging chamber with 5% CO_2_ using a Zeiss LSM980 microscope. 5 images from each condition were taken using 60X objective with Airyscan 2 in Multiplex mode.

Blinding of samples was not performed, however automated pipelines were used to quantify the data in an unbiased manner. Image analysis was performed using CellProfiler 4.1.3. Briefly, nuclei were identified with the ‘IdentifyPrimaryObjects’ module using global Otsu thresholding. Cellular boundaries were then identified with labelled nuclei as seeds, using propagation method within the ‘IdentifySecondaryObjects’ module. The outermost 10 pixels of each cellular boundary were saved as plasma membrane specific masks. Plasma membrane masks were then applied to each individual cells in Fiji (v2.1.0/1.53c).

Normalized generalized polarization ratio (GP ratio) of plasma membrane was quantified as described^20^.

Normalized GP ratio = (I_400-460_ – G*I_470-530_) / (I_400-460_ + G*I_470-530_), where

G = (GP_ref_ + GP_ref_*GP_mes_ – GP_mes_ – 1) / (GP_mes_ + GP_ref_*GP_mes_ – GP_ref_ – 1) GP_ref_ = 0.207^22^,

GP_mes_ was obtained from reference image with media containing the identical concentration of Laurdan (30μg/ml) without any cells. Reference images were taken in the same imaging session with identical parameters in all samples of the same experiment.

### *In vivo* Laurdan staining and quantification

Five young (3-5 month-old) and five old (20-22 month-old) C57BL/6 male mice were subjected to transcardiac perfusion with 4% paraformaldehyde (Electron Microscopy Sciences, 15714) in PBS (Corning, 21-040-CV). Brains were post-fixed overnight in 4% paraformaldehyde (Electron Microscopy Sciences, 15714) and then dehydrated in 30% sucrose (Sigma-Aldrich, S3929) for 72 hours. Brains were embedded in Tissue-Tek optimal cutting temperature (O.C.T.) compound (Electron Microscopy Sciences, 62550), sectioned into 16μm coronal slices using a cryostat (Leica CM3050S), and then mounted on glass slides (Fisher Scientific, 12-550-15). Coronal sections between Bregma 0.5mm to 1mm were generated for each mouse. Antibodies against Glial Fibrillary Acidic Protein (GFAP) (Abcam, ab53554) were used at 1:1000 dilution, antibodies against Epidermal Growth Factor Receptor (EGFR) (Millipore, 06-847) were used at 1:200 dilution at 4°C overnight. Donkey anti-goat antibodies conjugated with Alexa Fluor 647 (Thermo Fisher Scientific, A-21447), and donkey anti-rabbit antibodies with Alexa Fluor 568 (Thermo Fisher Scientific, A-10042) were used at 1:250 dilution at room temperature for 1.5 hours. Sections were subsequently stained with Laurdan at a final concentration of 30μg/ml for 30min at room temperature. Sections were then washed with PBS before being mounted with 50μl of Vectashield anti-fade mounting media (Vector Laboratories, H-1000-10) before being imaged.

Confocal imaging was carried out using a Zeiss LSM980 microscope using 60X objective with Airyscan 2 in Multiplex mode. Five to seven Z-stack images were taken for each individual animal with 10-15 images in each stack using 0.3μm intervals between images. Quiescent neural stem cells (qNSCs) were identified using the following criteria: 1) Positive GFAP staining surround the entire cell body in at least two consecutive images in the stack, 2) Positive GFAP staining partially detected throughout the rest of the images for the same cell, and 3) Negative EGFR staining throughout all images for the same cell. Once qNSCs were identified, plasma membrane-specific masks were hand-traced based on GFAP staining signal immediately surrounding the cell body and apical process using Fiji (v2.1.0/1.53c). Normalized generalized polarization ratio (GP ratio) of plasma membrane was quantified as described in the previous section.

### Fluorescence lifetime imaging (FLIM)

Membrane order was also assessed by fluorescence lifetime imaging by the polarity-sensitive membrane probe di-4-ANEPPDHQ (Invitrogen, D36802)^23^. Specifically, 90,000 qNSCs from primary culture of young and old mice were plated in each well of 8-well chamber coverslip (Ibidi, 80826). After 7 days of induced quiescence, cells were stained with di-4-ANEPPDHQ at a final concentration of 5μM in culture media for 1 hour in 37°C tissue culture incubator. FLIM imaging was carried out using a Zeiss LSM780 microscope using 60X objective. Excitation was set at 475 nm by a frequency-doubled, Mai Tai^®^ DeepSee^TM^ laser oscillator (Spectra-Physics). Fluorescence lifetimes were measured using a time-correlated single photon counting (TCSPC) card (Becker & Hickl GmbH) and fitted using an incomplete multiexponential model on the SPCM software (Becker & Hickl GmbH).

### CRISPR/Cas9 reagents

LentiCRISPR v2 was a gift from Feng Zhang (Addgene plasmid # 52961; http://n2t.net/addgene:52961; RRID:Addgene_52961). psPAX2 was a gift from Didier Trono (Addgene plasmid # 12260; http://n2t.net/addgene:12260; RRID:Addgene_12260). pCMV-VSV-G was a gift from Bob Weinberg (Addgene plasmid # 8454; http://n2t.net/addgene:8454; RRID:Addgene_8454). sgRNAs targeting (*Mboat2*, *Elovl5*, *Agpat3*, *Fads2*, *Pla2g4e*) together with non-targeting and safe-targeting sgRNAs were selected from previously published genome-wide sgRNA library^24^. For a complete list of sgRNAs, see Supplementary Table 7. sgRNA oligos were ordered from Integrated DNA Technologies and subcloned into lentiCRISPR v2 plasmid using the *BsmB*I restriction site as described^25^. Sanger sequencing was performed to verify the constructs. One sequencing primer verifying the insertion of sgRNA and the 5’-end of the Cas9 sequence was used in conjunction with a second sequencing primer verifying the 3’-end of the Cas9 sequence.

### Lentiviral production and transduction for CRISPR/Cas9-mediated knockouts

For lentiviral production, human embryonic kidney 293T cells were seeded at a density of 13×10^6^ cells per 15cm plate in DMEM (Thermo Fisher, 11965092) with 10% fetal bovine serum (Gibco, 10099-141, lot 1751896) and penicillin-streptomycin-glutamine 1X (Gibco, 10378-016). Two days after plating, 293T media was changed and the cells were transfected using 1mg/ml of polyethylenimine (PEI) (Polysciences, 23966-2).

LentiCRISPR v2 (45μg) was transfected together with the lentiviral packaging plasmids psPAX2 (28.35µg), and pCMV-VSV-G (8.325µg) per 15cm plate. One day (20-24 hours) after transfection, the media was changed to NeuroBasal-A with penicillin-streptomycin-glutamine. After another 20-24 hours, lentivirus-containing supernatant was collected and stored at 4°C and fresh media was added to the 293T cells for another collection after 24 hours. Both supernatants were then combined, filtered through a 0.45µm polyvinylidene fluoride (PVDF) filter (Millipore Sigma, SE1M003M00), and frozen at −80°C in 3ml aliquots. For lentiviral transduction, qNSCs were maintained in quiescence media for 5 days (with media change every other day) before transduction. After removing media, 750μl of lentiCRISPR v2 supernatant was thawed on ice and mixed with 250μl of NeuroBasal-A + penicillin-streptomycin-glutamine with 8% of B27 minus vitamin A, bFGF (80ng/ml) and BMP4 (200ng/ml). qNSC cultures were incubated with viral media for 24 hours. After removing viral media after 24 hours, a second lentiviral transduction was repeated the next day. After two consecutive transductions, cells were washed once with NeuroBasal-A media and then cells were kept in “quiescence NSC media” for 7 days to allow recovery and CRISPR editing. To assess the efficiency of viral transduction and reproducibility, a positive control lentiviral construct pLJM1-EGFP (Addgene, 19319) was used in parallel. The fraction of transduced cells was quantified as EGFP^+^ cells using an LSR-II flow cytometer (BD Biosciences) 48 hours after transduction. A transduction efficiency of 70-80% was observed throughout this study. Fresh quiescent NSC media was replaced every 2 days.

CRISPR editing efficiency was assessed by analyzing genomic DNA. Briefly, genomic DNA was collected in cells after 7 days of recovery period following lentiviral transduction. Genomic DNA was extracted after lysing cells with 200μl of DirectPCR lysis reagent (Viagen biotech, 102-T) for 15min at room temperature. Lysate was then digested with 0.2mg/ml Protease K (Invitrogen, 25530049) for 25min at 65°C, followed by protease deactivation for 15min at 95°C. We then amplified genomic DNA with primer pairs roughly 150-250bp upstream and 450-500bp downstream of sgRNA editing site, with 30s of annealing step at 55°C and 1min of extending step at 72°C for 40 cycles total. See Supplementary Table 8 for a complete list of primers.

We used the knockout score generated by Inference of CRISPR Edits (ICE) (Synthego Performance Analysis, ICE Analysis. 2019. v2.0. Synthego) to assess editing efficiency, according to the vendor’s instruction. Knockout score is calculated by ICE based on the fraction of alleles from the genomic DNA of edited samples with either a frameshift or 21+ bp indel, using an unedited sample as control. All knockouts performed in this study have an editing efficiency between 55-70%.

### Inference of CRISPR Edits (ICE) analysis for knockout efficiency

Genomic knockout efficiency was assessed using the ICE analysis tool (Synthego Performance Analysis, ICE Analysis. 2019. v3.0. Synthego; [2022]). Knockout score represents the percentage of alleles that contains a frameshift insertion/deletion in a given sample.

### Lipidomic aging score of quiescent NSCs with various knockouts

We used the set of lipidomic aging signatures (see above) to generate lipidomic aging score in assessing the effect of genetic knockout on global lipidome. We first calculated standard score (Z score) for each signature feature across all samples. The mean Z score of all signature features was then used as the lipidomic aging score for individual sample.

### NSC activation assay by FACS

To assess activation efficiency, qNSCs were washed twice with PBS (Corning, 21-040-CV) at 37°C before switching to “Proliferative NSC media” for 4 days. Media was replaced every 2 days. Cells were dissociated with Accutase (Stemcell Technologies, 07920) for 5min, collected into conical tubes, and centrifuged at 300g for 5min. Next, 1×10^6^ cells were resuspended in 500μl FACS buffer (PBS with 2% fetal bovine serum (Gibco, 10099-141, lot 1751896)). Following this step, 500μl of 3.2% formaldehyde, diluted from 16% formaldehyde (Thermo Scientific, 28906), was added to the suspension dropwise under agitation to achieve a final concentration of 1.6% formaldehyde. Cells were fixed in 1.6% formaldehyde for 10min at room temperature before spinning down at 700g for 5min. Cells were washed once with FACS buffer before permeabilizing with 1ml of ice-cold 100% methanol (Thermo Fisher Scientific, A4121).

Following a period of 20min incubation on ice, cells were then washed again and resuspended in 200μl of FACS buffer for staining. Ki67-APC (eBioscience, 17-5698-82) antibody was used at 1:300 dilution. Samples were incubated in the dark for 30min at room temperature with gentle agitation. Cells were washed again before adding DAPI (Thermo Scientific, 62248) at a final concentration of 0.2μg/ml in FACS buffer. Each sample was filtered with FACS-strainer cap tubes (Falcon, 352235), just prior to FACS analysis. Cells were analyzed on a BD LSR II flow cytometer, and Flowjo (v10) software was used for data analysis.

### Statistical analyses

We did not perform power analyses, though we did use previous experiments to inform the experimental design with respect to sample size. To calculate statistical significance for experiments, all tests were two-sided Wilcoxon rank-sum tests unless otherwise indicated. To test the effect of knockout in paired samples, paired Welch’s *t*-test was used. To test age difference within knockout condition, unpaired Welch’s *t*-test was used. Normality test was performed to ensure normal distribution before performing Welch’s *t*-test. All tests were performed by R version 4.0.2. All Results from individual experiments and all statistical analyses are included in Supplementary Table 9.

---

**Extended Data Figure 1.**
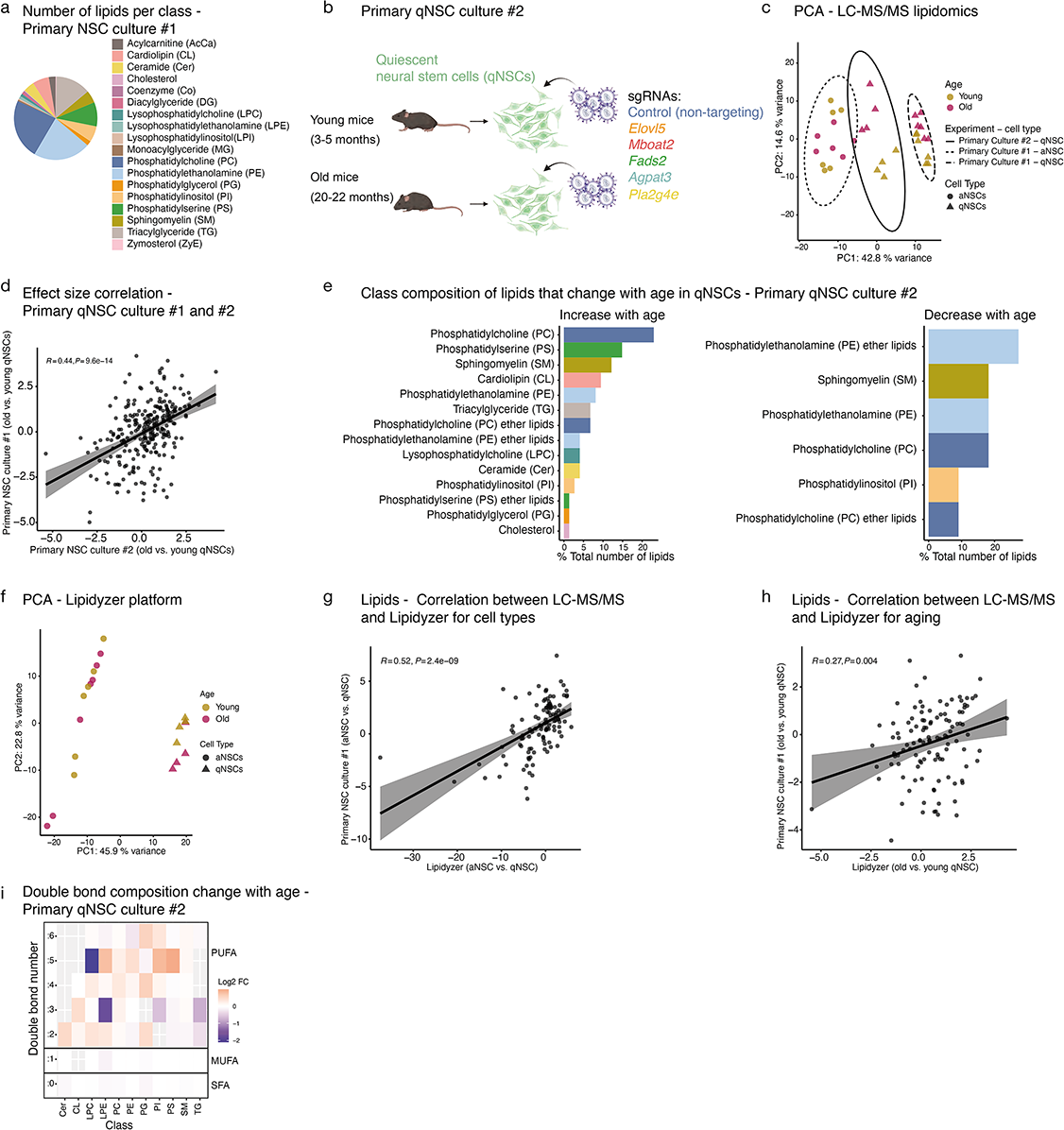
Reproducibility of lipidomic changes in primary neural stem cell cultures from independent experiments and platforms. **a,** Lipid composition of data from primary NSC culture #1 (qNSCs and aNSCs). Pie chart showing the number of lipids of each lipid class as a percentage to the total number of detected lipids. **b,** Samples included in primary qNSC culture #2. Primary culture of NSCs from young (3-5 months) and old (20-22 months) mice are induced into quiescence and then transduced with lentiviral-mediated CRISPR/Cas9 knockout containing a non-targeting sgRNA, or sgRNA targeting specific genes (lipidomics experiment also used in Figure 4). Samples with non-targeting control sgRNA were used as primary NSC culture #2 as an additional dataset to test the reproducibility of age-related lipidomic changes in qNSCs. **c,** Principal component analysis (PCA) with combined data from primary NSC culture #1 (aNSCs and qNSCs) and primary qNSC culture #2. Z score was obtained from log2-transformed data of each dataset. PCA was then generated using the common lipids that were detected in both datasets. Each point represents an individual primary aNSC cultures (dot) or qNSC cultures (triangle) established from young (golden) or old (pink) mice. A data ellipse is drawn at a 95% level for primary qNSC culture #2 (solid line), aNSCs of primary NSC culture #1 (dashed line) and qNSCs of primary NSC culture #1 (longdash line). **d,** Correlation between data from primary qNSC culture #1 and primary qNSC culture #2. Pearson correlation was performed on effect sizes between young and old qNSCs of individual lipids (black dots) calculated from each dataset. Linear regression line (black line) with 95% confidence interval (gray shade). **e,** Class composition of lipids that significantly change with age in qNSCs from primary NSC culture #2. Number of lipids in each lipid class as a percentage of the total number of lipids that show significant increase with age (left panel) or decrease with age (right panel). **f,** Principal component analysis (PCA) on lipids from Lipidyzer dataset. Log2-transformed normalized concentration of all lipids was used for this analysis. Each point represents an individual primary aNSC cultures (dot) or qNSC cultures (triangle) established from young (3-5 months, golden) and old (20-22 months, pink) C57BL/6JN male mice. **g, h,** Correlation between data from primary NSC culture #1 and targeted Lipidyzer platform. Pearson correlation was performed on effect sizes between aNSCs and qNSCs (**g**) and between young and old qNSCs (**h**) of individual lipids (black dots) calculated from each dataset. Linear regression line (black line) with 95% confidence interval (gray shade). **i,** Heatmap showing double bond composition change between with age in qNSCs from side chain composition analysis on primary qNSC culture #2. Log2 fold change in abundance of lipids with side chain containing between 0-6 double bonds (rows) in different classes (columns) are shown. Abbreviations defined in panel a.

**Extended Data Figure 2.**
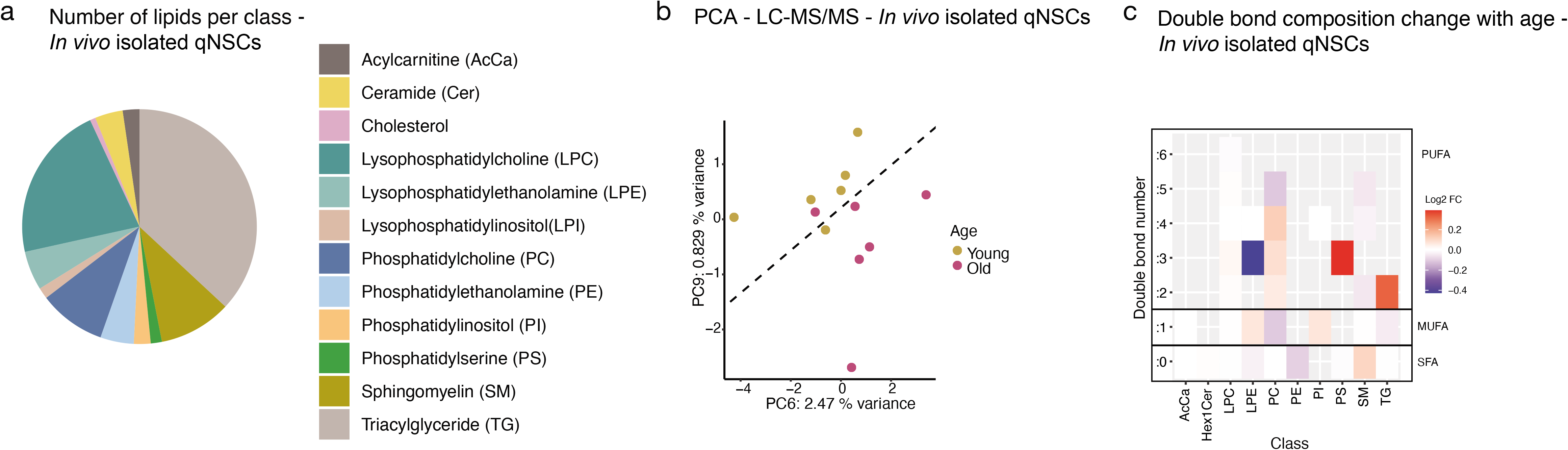
Lipidomics data from *in vivo* isolated quiescent neural stem cells from the subventricular zone neurogenic niche of young and old mice. **a,** Lipid composition of data from *in vivo* isolated qNSCs. Pie chart showing the number of lipids of each lipid class as a percentage to the total number of detected lipids. **b,** Principal component analysis (PCA) on log2-transformed normalized concentration or intensity of all lipids. Specifically, for lipids that have internal standard for their respective lipid classes, molar concentration was used, for lipid classes that don’t have internal standard, normalized intensity was used in the PCA analysis. Each point represents an individual young (3-5 months, golden) or old (20-22 months, pink) C57BL/6JN male mouse. A computed decision boundary indicating the separation between young and old samples (see Methods) was added to the PCA plot (dashed line). **c,** Heatmap showing double bond composition change with age in qNSCs from side chain composition analysis on *in vivo* isolated qNSCs. Log2 fold change in abundance of lipids with side chain containing between 0-6 double bonds (rows) in different classes (columns) are shown. Abbreviations defined in panel a, except for Hex1Cer: hexosylceramides.

**Extended Data Figure 3.**
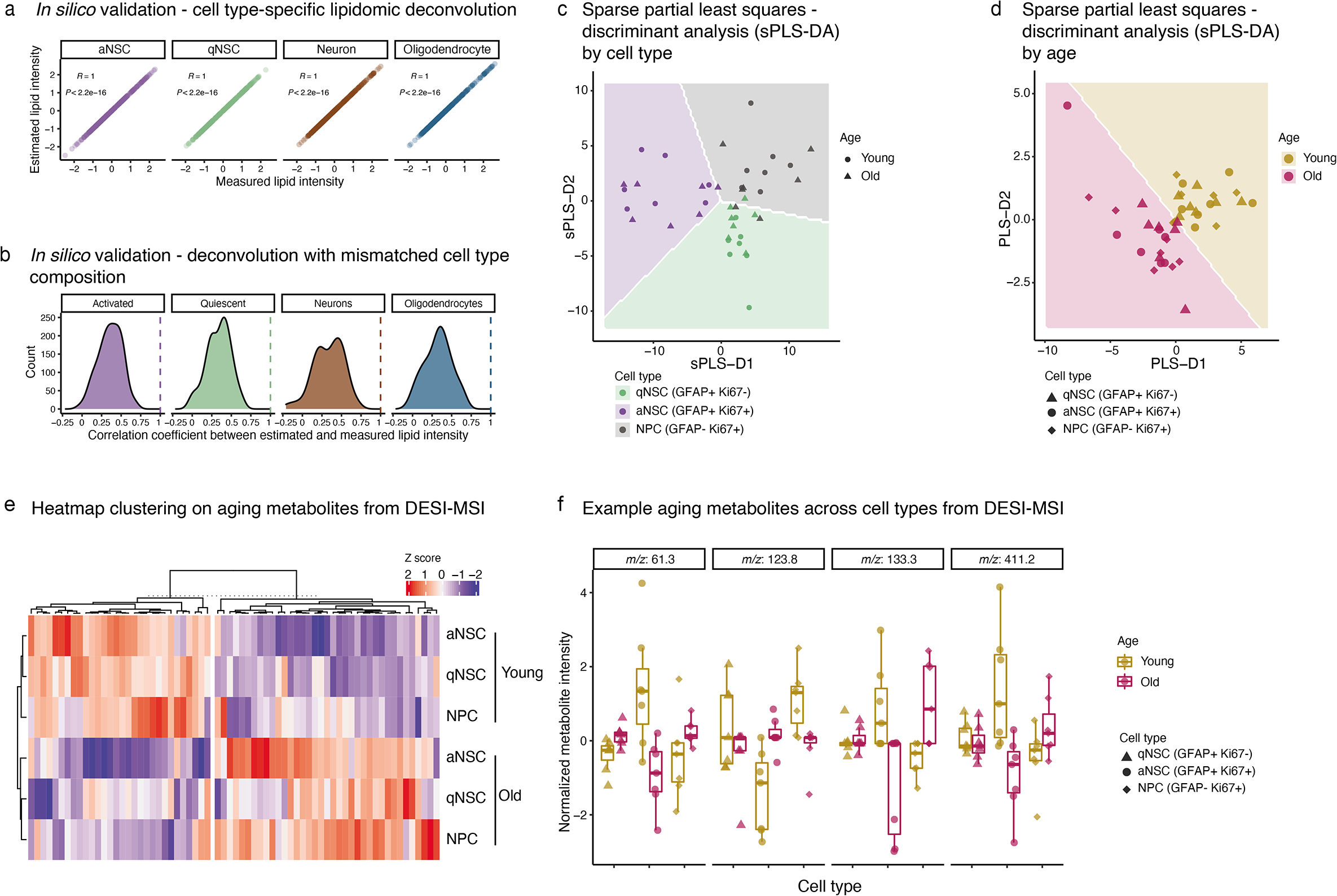
Metabolomic profiling on young and old subventricular zone neurogenic niches by DESI-MSI. **a,** Cell type-specific deconvolution from *in silico* mixture lipidomic data. Pearson correlation was performed between the Z score of measured lipid intensity and Z score of estimated lipid intensity of activated NSC (aNSC), quiescent NSC (qNSC), neurons and oligodendrocytes from a reconstituted mixture lipidomic dataset (see Methods). Each dot represents a lipid species detected from an individual culture of brain cell type. **b,** Deconvolution from *in silico* mixture lipidomic data with mismatched cell type proportions using randomly-generated cell type compositions (see Methods). Density plot shows the number of occurrence (Y axis) of the Pearson correlation coefficient (X axis) of each lipid species between measured and estimated lipid intensity for each individual cell type after deconvolution. Dashed line indicates the correlation coefficient obtained from deconvolution when the matching cell proportions were used (same as panel a). **c, d,** Sparse partial least squares-discriminant analysis (sPLS-DA) on DESI-MSI metabolomic profiling of SVZ cells from 7 young (3-5 months) and 7 old (20-22 months) C57BL/6JN male mice. sPLS-DA was performed to obtain cell type-specific (**c**) and age-specific (**d**) metabolic signatures. Each data point represents a deconvolved metabolomic profile of a specific cell type from an individual young or old mouse. Cell types defined by immunostaining for GFAP (NSC, astrocytes) and Ki67 (proliferation). aNSC: GFAP+Ki67+; qNSC: GFAP+Ki67-; NPC: GFAP-Ki67+. **e,** Heatmap clustering of metabolite features associated with aging differences across 3 cell types in the SVZ neurogenic niche (see Methods). Z score on the intensity of each metabolite (columns) in different cell types (rows) are shown. **f,** Metabolites with unique cell type-specific changes with age. Metabolites were selected based on significant interaction between age and cell type by ANOVA analysis. Each data point represents the normalized peak intensity of a given metabolite in different SVZ cell types of one individual mouse. Cell types defined by immunostaining for GFAP (NSC, astrocytes) and Ki67 (proliferation). qNSC: GFAP+Ki67-; aNSC: GFAP+Ki67+; NPC: GFAP-Ki67+.

**Extended Data Figure 4.**
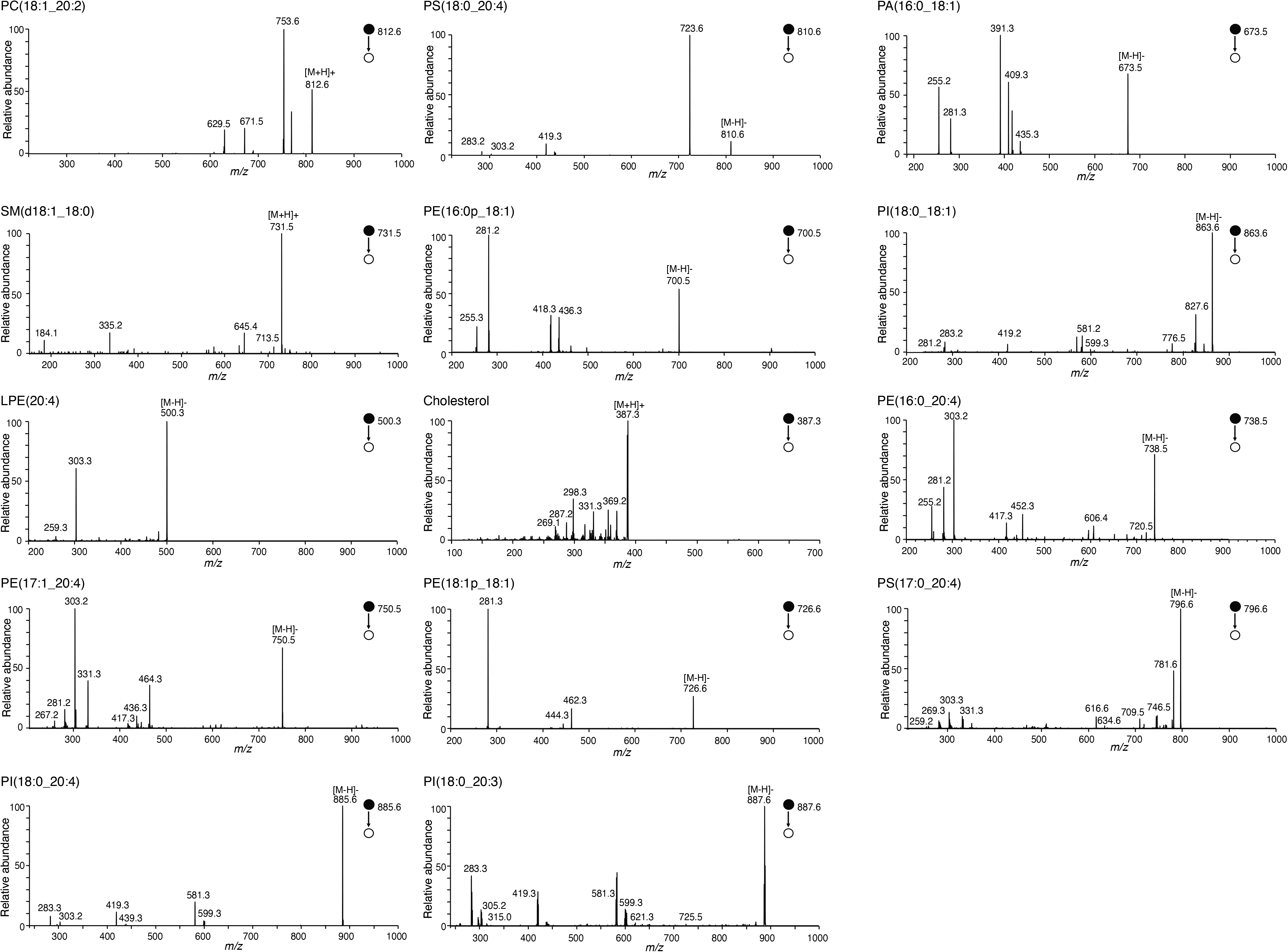
Tandem mass spectrometry validation following DESI-MSI. Structural validation by tandem mass spectrometry for PC(18:1_20:2), PS(18:0_20:4), PA(16:0_18:1), SM(d18:1_18:0), PE(16:0p_18:1), PI(18:0_18:1), LPE(20:4), Cholesterol, PE(16:0_20:4), PE(17:1_20:4), PE(18:1p_18:1), PS(17:0_20:4), PI(18:0_20:4), and PI(18:0_20:3). Solid dot in all spectra represents molecular mass used to identify each molecule (precursor ion).

**Extended Data Figure 5.**
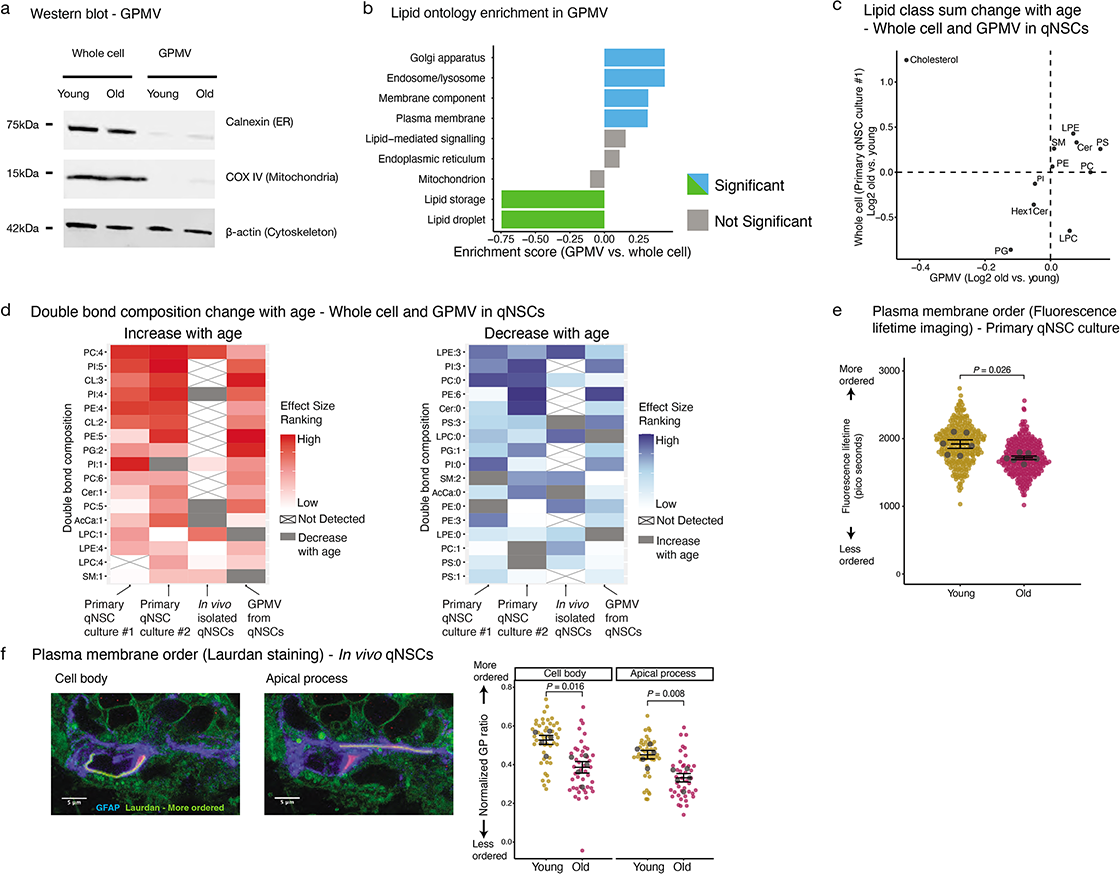
Plasma membrane lipidomic and membrane order change with age in quiescent neural stem cells. **a,** Western blot on protein extracts from whole cell lysates or giant plasma membrane vesicles (GPMVs) generated from primary qNSC cultures from young (3-5 months) and old (20-22 months) mice. Membranes were incubated with antibodies to cytoskeleton marker β-actin, endoplasmic reticulum marker Calnexin, and mitochondria marker COX IV. **b,** Lipid ontology analysis on lipids of different cellular components between GPMV lipidomics and whole cell lipidomics (primary qNSC culture #1). Enrichment score of each cellular compartment is presented. Terms with positive enrichment score are enriched in GPMV lipids, while terms with negative enrichment score are enriched in lipids from whole cell extract. Significant terms (FDR<0.05) are shown in blue/green color. Terms without statistical significance are shown in gray. **c,** Lipid class abundance change with age in whole cell (primary qNSC culture #1) and GPMV lipidomic data. Dot plot showing the log2 fold change with age on the total intensity or total concentration of each lipid class from whole cell and GPMVs generated from primary qNSC cultures of young and old mice. **d,** Double bond composition change with age in primary qNSC cultures, *in vivo* qNSCs, and GPMVs from qNSCs. Aging effect size of double bond composition was calculated, and then used to generate a percentile ranking for each dataset. Consistent double bond composition features that are detected in at least 3 out of the 4 datasets that show increase with age (left panel) or decrease with age (right panel) are then sorted based on the combined percentile ranking across all 4 studies. Heatmap is plotted based on the percentile ranking of double bond features in each lipidomic dataset. Double bond features not detected is crossed out. Gray indicates double bond features with opposite age-related changes from other datasets. **e,** Plasma membrane order assessed *in vitro* by fluorescence lifetime imaging (FLIM) on primary qNSC cultures established from young (3-5 months) and old (20-22 months) mice. SuperPlots showing data from individual cells (golden/pink dots), as well as the mean of each primary culture from individual mice (gray dots) +/-SEM. Results from two independent experiments combined, n=6 primary cultures. *P*-value from Wilcoxon rank-sum test. **f,** Plasma membrane lipid order assessed by Laurdan of different subcellular regions in qNSCs *in vivo* from 5 young (3-5 months) and 5 old (20-22 months) mice. Left: representative Laurdan staining on *in vivo* qNSCs from SVZ. Image taken from rigid wavelength (more ordered, green) is overlaid with GFAP (NSC and astrocyte marker, blue). Plasma membrane segment around the cell body as well as around the apical process is outlined in yellow. Right: Membrane order quantified by generalized polarization ratio (GP ratio) from Laurdan staining. SuperPlots showing data from individual cells (golden/maroon dots), as well as the mean of each mouse (gray dots) +/-SEM. Results are from one independent experiment, n=5 mice. *P*-value from Wilcoxon rank-sum test. Results from a second independent experiment are in Supplementary Table 9.

**Extended Data Figure 6.**
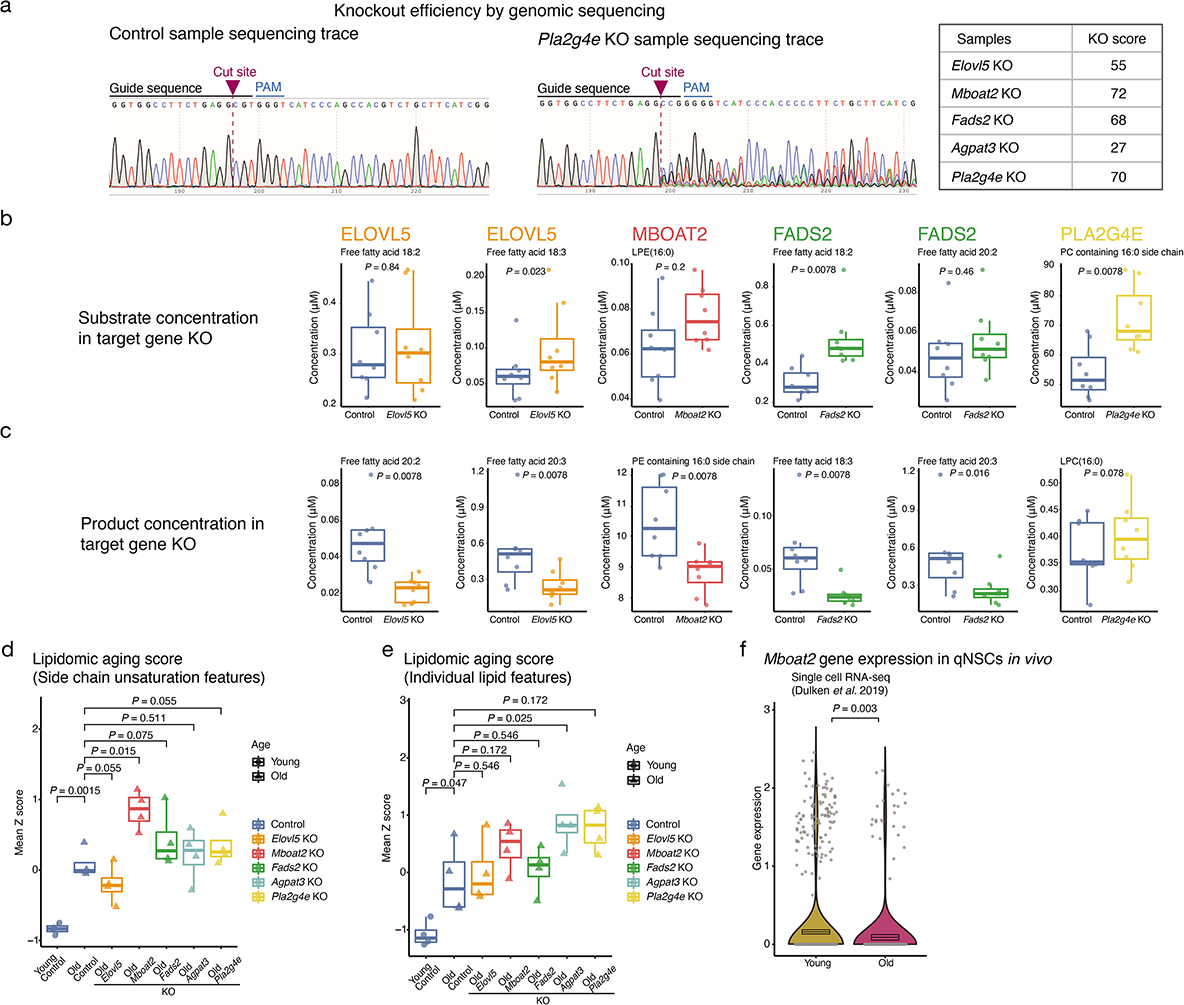
Validation of gene knockouts and effect on the global lipidome of neural stem cells. **a,** Validation of knockout efficiency by genomic DNA sequencing. Representative sequencing traces from control and knockout (KO) samples of the *Pla2g4e* gene with CRISPR editing are shown (left). Summary of KO score is obtained from Inference of CRISPR Edits (ICE) analysis (see Methods) (right). **b, c,** Substrate (b) and product (c) level in control and knockout (KO) for each target genes in qNSCs. Data is obtained from untargeted lipidomic analysis. Each dot represents a primary culture from an individual young (3-5 months old) or old (20-22 months old) male mouse. Results are from 1 independent experiment, n=8 primary cultures (4 young and 4 old). *P*-values from Wilcoxon rank-sum test. Substrates are: *Elovl5* KO: free fatty acid 18:2 and 18:3, *Mboat2* KO: LPE(16:0), *Fads2* KO: free fatty acid 18:2 and 20:2 and *Pla2g4e* KO: PC containing 16:0 side chain. Product are: *Elovl5* KO: free fatty acid 20:2 and 20:3, *Mboat2* KO: PE containing 16:0 side chain, *Fads2* KO: free fatty acid 18:3 and 20:3 and *Pla2g4e* KO: LPC(16:0). **d, e,** Lipidomic aging score calculated using side chain unsaturation aging signatures (**d**) and individual lipid aging signatures (**e**) from young (blue, circle) and old control samples (blue, triangle) and old samples with *Elovl5* knockout (KO) (orange, triangle), *Mboat2* KO (red, triangle), *Fads2* KO (green, triangle), *Agpat3* KO (turquoise, triangle) or *Pla2g4e* KO (yellow, triangle). Each data point represents a primary culture from an individual mouse. Results are from 1 independent experiment, n=4 primary cultures. *P*-value between old KO samples were from paired Welch’s *t*-test with multiple hypothesis correction by FDR. Normality of the data was confirmed for use of *t*-test. *P*-value between old and young control samples were from unpaired Welch’s *t*-test (young and old samples were not paired). Normality of the data was confirmed for use of *t*-test. **f,** Violin plots of *Mboat2* gene expression in young and old qNSCs/astrocytes from single-cell RNA-seq datasets. Boxplot: mean expression level +/-SEM. Each individual dot represents the normalized gene expression of *Mboat2* in a single cell. Gray bar at the bottom represents cells with gene expression level below detection limit. *P*-value from Wilcoxon rank-sum test.

**Extended Data Figure 7.**
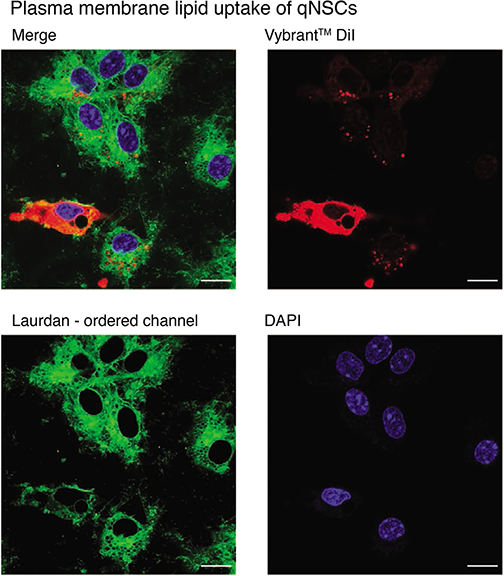
Uptake of lipids in quiescent neural stem cells upon plasma lipid supplementation. Representative confocal fluorescence images of primary qNSC cultures after plasma membrane lipid supplementation for 72 hours. Cells were imaged with Laurdan rigid wavelength (green), lipophilic dye Vybrant^TM^ DiI for labelling lipid extract (red), and DAPI (blue). Scale bar: 10μm. Labelled supplemented lipids were found in puncta, indicating successful uptake by the cell through the endocytic system. Endocytosed lipids can then be used by different compartments, including plasma membranes. Excess intracellular Vybrant^TM^ DiI can diffuse laterally and stain the entire cell.

**Extended Data Figure 8.**
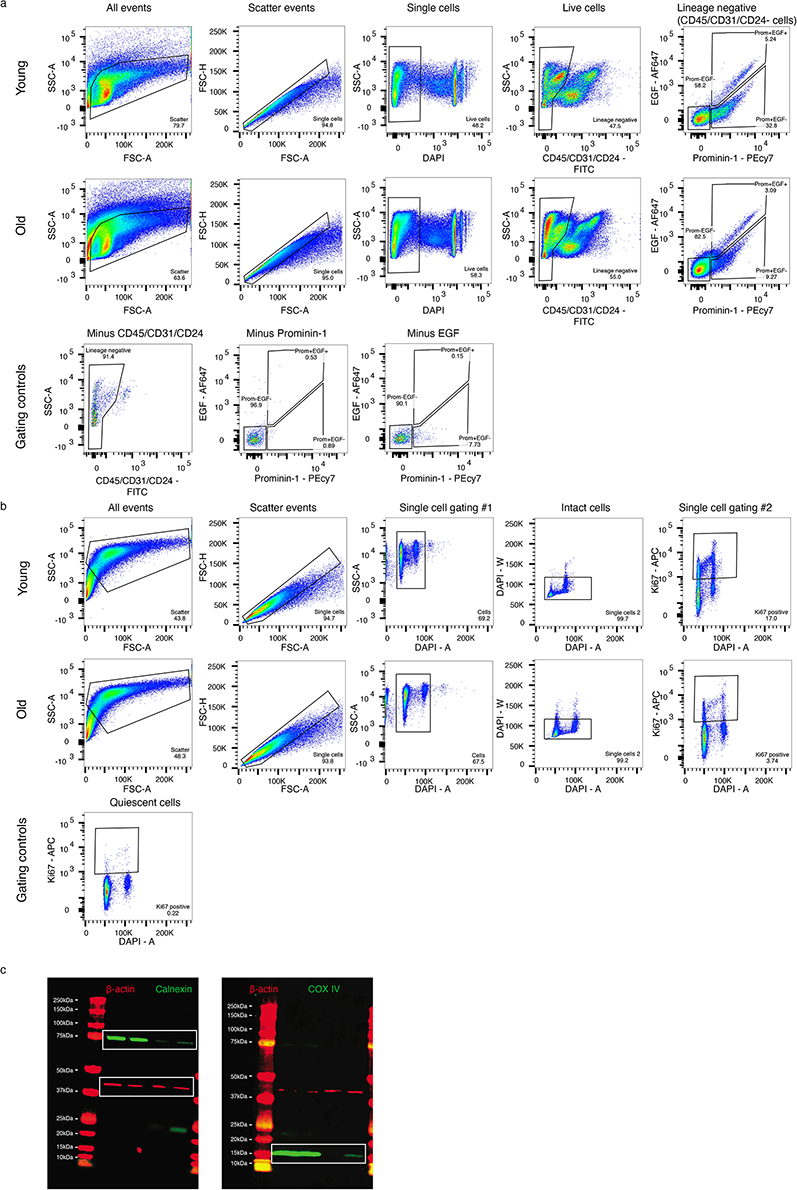
FACS gating scheme and full western blot membranes. **a,** FACS gating strategy used to freshly isolate quiescent NSCs from the SVZ neurogenic niches of young and old mice. Negative gating controls for each staining are indicated. For gating controls, cells were stained with all antibodies except the one for which it is a negative control. **b,** FACS gating strategy to assess the ability of quiescent NSCs to activate. Quiescent (non-proliferating) NSCs were used as negative control for the proliferation marker Ki67. The cell population on which each respective FACS gating was done is indicated above each plot. **c,** Full western blot membranes from Extended Data Fig. 5a. Boxes indicate the cropped areas used in Extended Data Fig. 5a (after conversion to gray scale).

## Supplementary Tables

Supplementary Table 1: Lipidomic data – Young and old primary NSC culture #1

Supplementary Table 2: Lipidomic data – Young and old primary qNSC culture #2

Supplementary Table 3: Lipidomic data – Young and old primary NSC culture by Lipidyzer

Supplementary Table 4: Lipidomic data – Young and old *in vivo* isolated qNSCs

Supplementary Table 5: Lipidomic data – Young and old *in situ* DESI-MSI in neurogenic niches

Supplementary Table 6: Lipidomic data – Young and old GPMVs from primary qNSC culture

Supplementary Table 7: List of sgRNAs for knockout

Supplementary Table 8: List of sequencing primers

Supplementary Table 9: Experimental data and statistics

## Notes

### Competing Interest Statement

The authors have declared no competing interest.

